# Acarbose Impairs Gut *Bacteroides* Growth by Targeting Intracellular GH97 Enzymes

**DOI:** 10.1101/2024.05.20.595031

**Authors:** Haley A. Brown, Adeline L. Morris, Nicholas A. Pudlo, Ashley E. Hopkins, Eric C. Martens, Jonathan L. Golob, Nicole M. Koropatkin

## Abstract

Acarbose is a type-2 diabetes medicine that inhibits dietary starch breakdown into glucose by inhibiting host amylase and glucosidase enzymes. Numerous gut species in the *Bacteroides* genus enzymatically break down starch and change in relative abundance within the gut microbiome in acarbose-treated individuals. To mechanistically explain this observation, we used two model starch-degrading *Bacteroides*, *Bacteroides ovatus* (Bo) and *Bacteroides thetaiotaomicron* (Bt). Bt growth is severely impaired by acarbose whereas Bo growth is not. The *Bacteroides* use a starch utilization system (Sus) to grow on starch. We hypothesized that Bo and Bt Sus enzymes are differentially inhibited by acarbose. Instead, we discovered that although acarbose primarily targets the Sus periplasmic GH97 enzymes in both organisms, the drug affects starch processing at multiple other points. Acarbose competes for transport through the Sus beta-barrel proteins and binds to the Sus transcriptional regulators. Further, Bo expresses a non-Sus GH97 (BoGH97D) when grown in starch with acarbose. The Bt homolog, BtGH97H, is not expressed in the same conditions, nor can overexpression of BoGH97D complement the Bt growth inhibition in the presence of acarbose. This work informs us about unexpected complexities of Sus function and regulation in *Bacteroides*, including variation between related species. Further, this indicates that the gut microbiome may be a source of variable response to acarbose treatment for diabetes.

**Importance:** Acarbose is a type 2 diabetes medication that works primarily by stopping starch breakdown into glucose in the small intestine. This is accomplished by inhibition of host enzymes, leading to better blood sugar control via reduced ability to derive glucose from dietary starches. The drug and undigested starch travel to the large intestine where acarbose interferes with the ability of some bacteria to grow on starch. However, little is known about how gut bacteria interact with acarbose, including microbes that can use starch as a carbon source. Here, we show that two gut species, *Bacteroides ovatus* (Bo) and *Bacteroides thetaiotaomicron* (Bt), respond differently to acarbose: Bt growth is inhibited by acarbose while Bo growth is not. We reveal a complex set of mechanisms involving differences in starch import and sensing behind the different Bo and Bt responses. This indicates the gut microbiome may be a source of variable response to acarbose treatment for diabetes via complex mechanisms in common gut microbes.

## Introduction

Recent work suggests that the beneficial or toxic effects of some xenobiotics are mediated via their interactions with the gut microbiota (1). For example, the severe negative side effects of the colon cancer chemotherapeutic, CPT-11, are caused by enzymatic reactivation of the drug by symbiotic intestinal bacteria (2). Conversely, metformin, which is used to treat type 2 diabetes (T2D), elicits some of its positive effects due to a direct influence on the composition of bacteria in the gut (3). The natural product acarbose (**Fig. 1A**), is also FDA approved to treat T2D. Though the FDA has long known that acarbose “…is metabolized exclusively within the gastrointestinal tract, principally by intestinal bacteria,” there is a paucity of data providing molecular insight into these interactions (4). A mechanistic investigation into acarbose bacterial interactions is therefore warranted.

**Figure 1.**
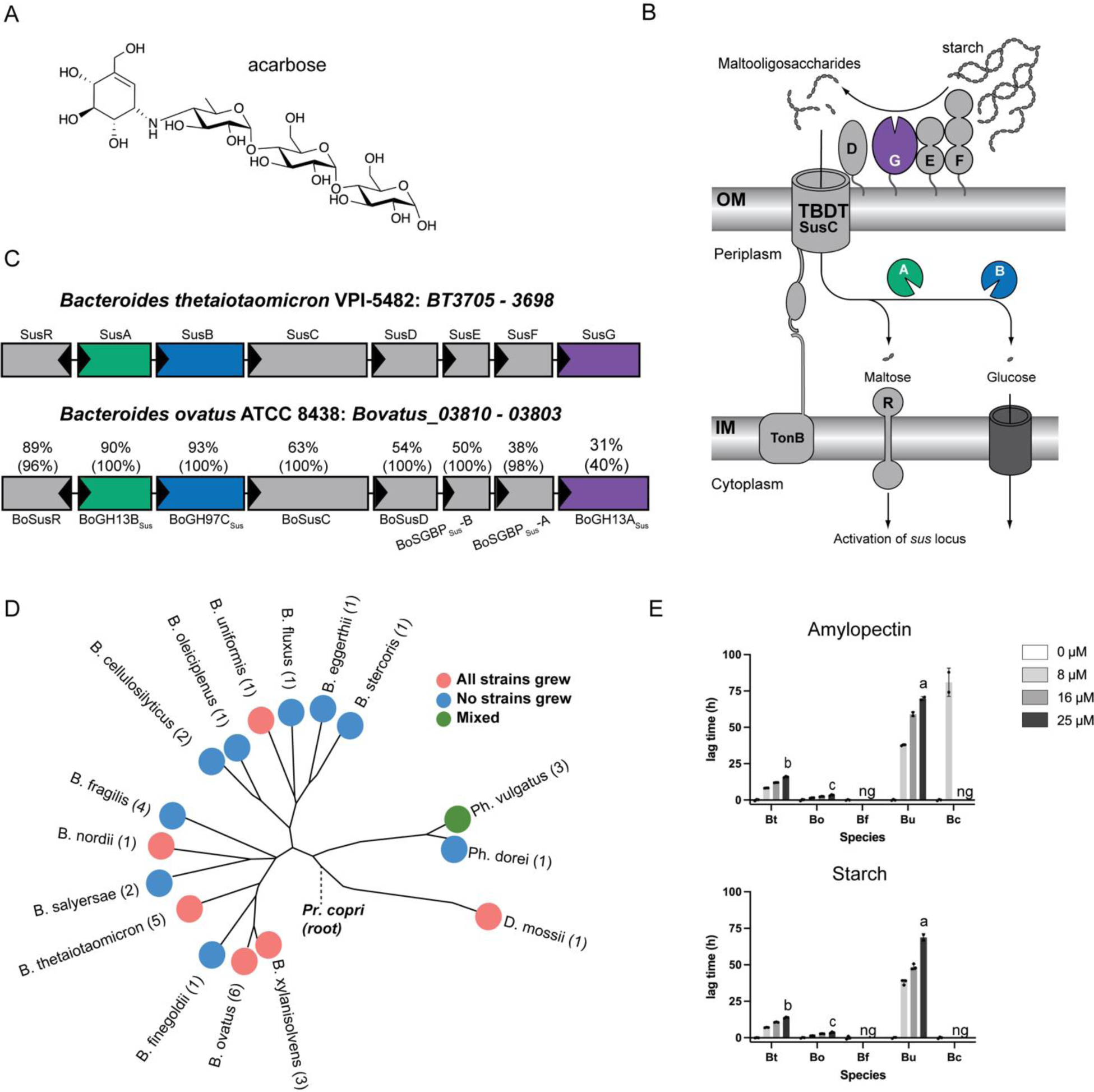
*Bacteroides* growth is inhibited by acarbose. A) Chemical structure of acarbose. B) Summary of the Bt starch utilization system with an emphasis on the likely enzymatic targets of acarbose: the outer member GH13 SusG, periplasmic GH97 SusB, and periplasmic GH13 SusA. C) Amino acid level comparison of Bt and Bo Sus. Numbers above the Bo proteins correspond to percent identity and sequence coverage. For example, BoSusC is 63% identical to SusC over 100% of the sequence. B and C were modified from (27) and (51). D) Various *Bacteroides* strains and species were grown in amylopectin or potato starch with or without 25 µM acarbose. Growth was defined as reaching an OD_600_ of 0.3 over 144 hours of measured growth. The 16s rRNA based phylogenetic tree was modified from (52). E) Type strains of Bt, Bo, *Bacteroides fragilis* (Bf)*, Bacteroides uniformis* (Bu), and *Bacteroides cellulosilyticus* (Bc) were grown in the indicated concentrations of acarbose in amylopectin and potato starch. Lag time is defined as the difference in time it takes the treated and untreated conditions to reach an OD_600_ of 0.3. Statistical comparisons were performed for the 25 µM treatment only using Bo as the comparison point. P-values using a one-way ANOVA analysis were calculated. All comparisons were p<0.05, ng = no growth. All species grew in the 0 µM acarbose treatment.

The direct function of acarbose is to prevent postprandial blood glucose spikes by inhibiting host amylases and glucosidases (collectively termed glucoamylases) used to digest starch in the upper gastrointestinal tract (5). Acarbose functions as a transition state mimic for α1,4 glycosidic bond hydrolysis to inhibit these enzymes (6). During acarbose treatment, more starch transits to the large intestine where it is bacterially fermented to short chain fatty acids (SCFAs), leading to increased output of the beneficial SCFA, butyrate (7–9). Acarbose also changes the microbially derived bile acid pool. This is thought to contribute to its anti-hypoglycemic effects by influencing host metabolic signaling (10, 11).

In addition to being an effective treatment for T2D, acarbose shows promise in ameliorating symptoms in immune and cardiovascular diseases and consistently enhances murine lifespan, likely due to altered SCFA output and the interplay between glucose metabolism and aging (12–17). The drug was the subject of two clinical trials because of its anti-aging phenotype in mice and two for its promise in treating other diseases (18–21). Because acarbose efficacy may depend on an individual’s baseline microbiota and genetic predisposition to disease (10), molecular insight into the drug’s influence on bacterial growth and fitness is critical to repurposing this drug and improving outcomes in diseases besides T2D.

Since acarbose is minimally absorbed by host tissue and transits the gut, one explanation for its gut microbial effects is that it changes bacterial fitness by inhibiting bacterial glucoamylases (22, 23). Indeed, preliminary work shows that gut bacterial species are differentially inhibited by acarbose *in vitro* when grown in starch (24–26). In particular, members of the prevalent Bacteroidota phylum may be less fit since their relative abundance decreases in acarbose treated individuals (10). Species in this phylum normally deploy a well-characterized <u>S</u>tarch <u>u</u>tilization <u>s</u>ystem (Sus) to recognize, degrade, and import starch (**Fig. 1B**) (27). Starch is comprised of amylose and amylopectin. Amylose is made of almost exclusively α1,4-linked glucose while amylopectin contains α1,6 branch points (28). In the model organism *Bacteroides thetaiotaomicron* (Bt), Sus is comprised of the outer membrane lipoproteins SusDEF that bind starch and a lipoprotein α-amylase, SusG, that hydrolyzes substrate (29–38). Maltooligosaccharides are transported to the periplasm by a predicted TonB-dependent transporter, SusC, where they are degraded to glucose by a neo-pullulanase/α-amylase, SusA, and an α-glucosidase/glucoamylase, SusB (39–43). SusR senses periplasmic maltose and upregulates *sus* transcription (44).

Some gut isolates lacking a pathway analogous to Sus encode an acarbose kinase and acarbose glucosidase that inactivate the drug (45, 46). Still other enzymes from non-gut isolates can hydrolyze acarbose (47–49). The *Bacteroides* do not have homologues to these enzymes which made us question how or if they can circumvent acarbose induced growth inhibition. Recent work demonstrated that some *Bacteroides* can take up fluorescently labelled maltodextrin in the presence of acarbose even while acarbose impairs growth (26). While this suggests acarbose can target an intracellular enzyme, this has not been shown mechanistically.

Bt and another model member of the Bacteroidota phylum, *Bacteroides ovatus* (Bo) have nearly identical *sus* loci (**Fig 1C**). Strikingly, Bt growth on starch is inhibited by acarbose, but Bo is resistant to this growth inhibition. *In vitro* experiments demonstrate that growth on glucose and some non-starch substrates in acarbose is normal, strongly suggesting that one or more molecular features of Sus is responsible for the different phenotypes (50). The extracellular α-amylase is the least conserved protein between the two organisms with only 31% identity and 40% coverage (51) (**Fig 1C**). Since acarbose inhibits glucoamylases, we initially hypothesized that BoGH13A_Sus_ and SusG were differentially inhibited by acarbose. Via genetic manipulation, growth experiments, and inhibition kinetics of all Bo and Bt Sus enzymes, here we show that although the primary target of acarbose in Bo and Bt is their respective periplasmic GH97 Sus enzyme, these enzymes do not explain the drastically different Bo and Bt acarbose phenotypes. To our surprise, Bo upregulates a non-Sus α-glucosidase gene (*bovatus_04772*), here termed BoGH97D, when exposed to maltose or acarbose. Bt encodes a nearly identical protein, BtGH97H, but its gene is not upregulated in conditions where acarbose impairs growth. Furthermore, constitutive expression of BoGH97D in Bt does not rescue its acarbose induced lag phenotype. Instead, other non-enzymatic Sus features such as the TonB-dependent transporter and periplasmic transcriptional regulator may underpin Bo’s resistance to acarbose induced growth inhibition.

Our work underscores the importance of interrogating the molecular interactions of acarbose with individual bacteria and the prominent role of GH97 enzymes in driving the observed acarbose phenotypes. Furthermore, we revealed that acarbose interferes not only with enzymatic starch breakdown, but the ability of bacteria to transport and sense starch. Our data provide mechanistic insight into why members of the Bacteroidota may decrease in relative abundance in acarbose treated individuals and influence how patients respond to acarbose treatment.

## Results

### Bacteroides thetaiotaomicron is more sensitive to growth inhibition by acarbose than Bacteroides ovatus

*Bacteroides* growth inhibition by acarbose has been previously described for *Bacteroides thetaiotaomicron* (Bt), *Bacteroides fragilis* (Bf), *Phocaeicola dorei*, *Phocaeicola vulgatus,* and *B. xylanisolvens* (24–26). We extended these data and screened various Bacteroidota species for growth in potato starch and amylopectin with or without 25 µM acarbose (**Fig. 1D**). All strains of a given species either grew or didn’t grow in both polysaccharides with acarbose after 144 hours except *Phocaeicola vulgatus* which was mixed. Within species that did grow, we observed a range of acarbose induced lags (difference in time to an OD_600_ of 0.3 in acarbose vs. untreated control) (**Fig. 1E**). Bf was inhibited in as low as 8 µM acarbose whereas *Bacteroides cellulosilyticus* (Bc) grew in amylopectin with 8 µM acarbose but was otherwise inhibited. *Bacteroides uniformis* (Bu) grew in 25 µM acarbose with a very extended lag compared to Bt and *Bacteroides ovatus* (Bo). Given that Bo and Bt are both genetically tractable but exhibit different acarbose induced lag times, we used these species as models to determine how acarbose inhibits bacterial growth.

The Bt <u>s</u>tarch <u>u</u>tilization <u>s</u>ystem (Sus) is one model to study polysaccharide utilization loci (PUL) function in the Bacteroidota. Although BoSus has not been as extensively characterized, the organization of its *sus* locus is nearly identical to that of Bt (**Fig. 1B,C**). We were therefore surprised that Bo did not respond to acarbose similarly to Bt (**Fig. 1E**, **Fig. 2A**). In a previous study, 10 μM acarbose led to nearly complete inhibition of Bt growth after 18 hours on pullulan, a fungal polysaccharide comprised of repeating α1,6 linked maltotriose units, and potato starch (24). We repeated these experiments and extended the acarbose concentrations and polysaccharides tested.

**Figure 2.**
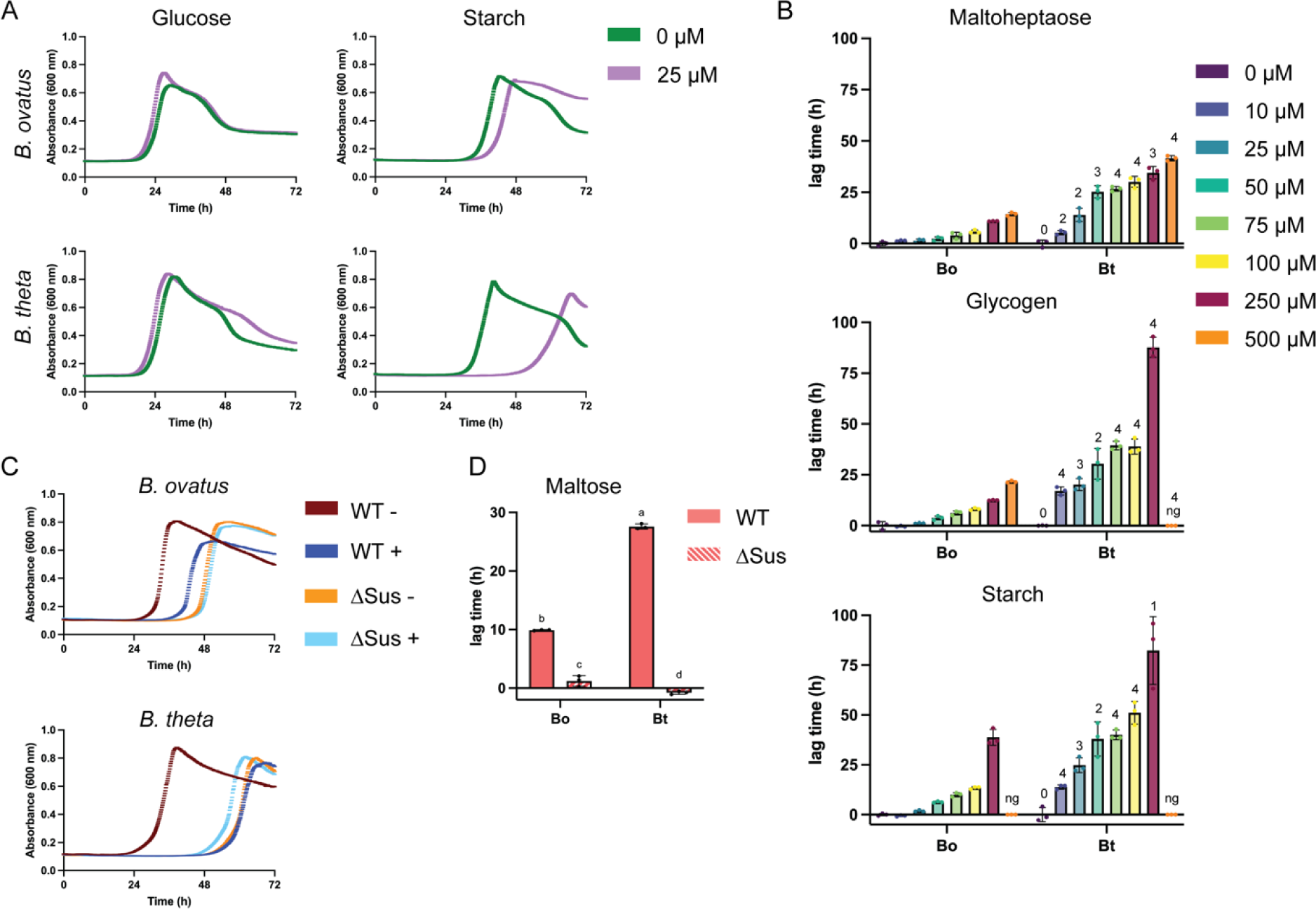
Bt is more susceptible to acarbose induced growth inhibition than Bo. A) Bo and Bt were pre-grown in minimal media (MM) with glucose and back diluted into MM + 2.5 mg/ml of the indicated carbon sources +/− 25 µM acarbose. B) Bo and Bt were pre-grown as described in A and back diluted into MM + 2.5 mg/ml of the indicated carbon sources with and without 10 – 500 µM acarbose. The difference in time to OD_600_ of 0.3 between the treated and untreated conditions are graphed. Bo and Bt with the same treatment were compared using an unpaired, two-tailed Student’s *t* test. 1: p ≤ 0.05; 2: p ≤ 0.01; 3: p ≤ 0.001. 4: p ≤ 0.0001. ng = no growth. C) WT and ΔSus strains of Bo and Bt were pre-grown as described in A and back diluted into MM + 2.5 mg/ml maltose +/− 50 µM acarbose. - = no acarbose and + = 50 µM acarbose. D) Lag time as described in B is shown from curves in C). Statistical analyses were performed using a two-way ANOVA with a cutoff of p≤0.05. Condition(s) with the same letter were not statistically different from one another.

To quantitatively assess the effects of acarbose on bacterial growth, we grew Bo and Bt in minimal media (MM) + 2.5 mg/ml glucose, maltose, maltoheptaose, glycogen, pullulan, amylopectin, and starch alone or with 10 – 500 µM acarbose (**Fig. 2B** and **Fig. S1A**). From this we calculated an IC_50_, which we defined as half the lowest acarbose dose that exhibited growth in at least two replicates by 72 hours (**Table 1**). Both Bo and Bt tolerate up to 500 µM acarbose in all oligosaccharides tested: glucose, maltose, and maltoheptaose (G7). When grown in pullulan, glycogen (more α1,6 branch points than amylopectin), amylopectin, or starch, Bt tolerates much less acarbose than Bo, except in starch (**Table 1**) (28). However, these IC_50_s do not capture the striking differences in acarbose induced lag times between Bo and Bt. At every concentration tested and with all tested α-glucans, Bt exhibited a significantly longer lag time than Bo (**Fig. 2B, Fig. S1A**).

**Table 1.**
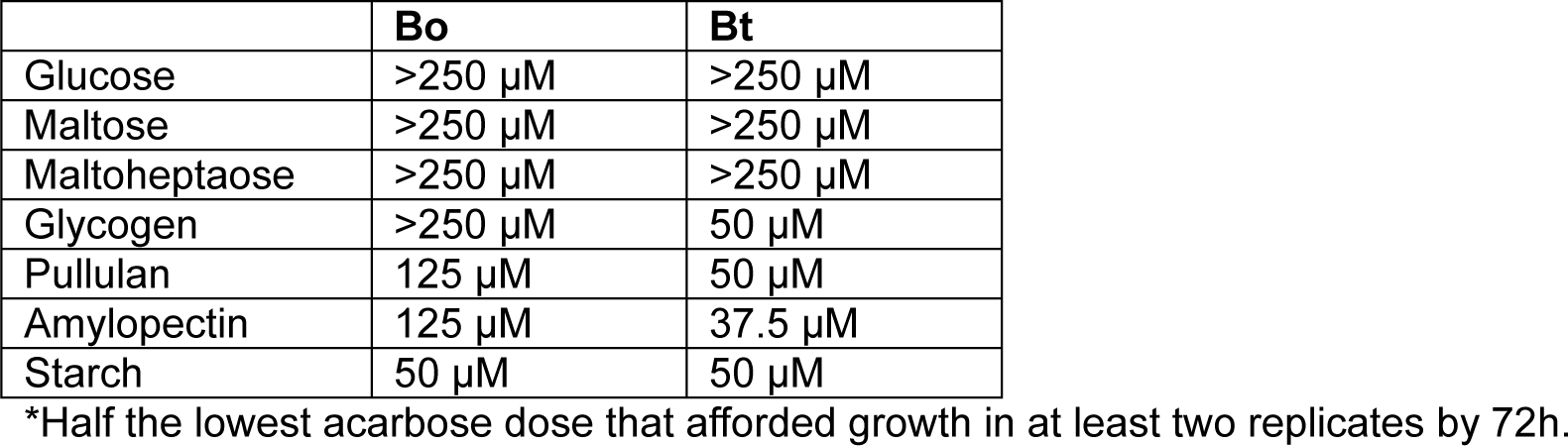
Acarbose IC_50_ values*.

The difference in acarbose tolerance does not appear to be due to breakdown of the drug, as cell lysates of Bo and Bt grown on maltose cannot break down acarbose, even after an extended incubation (**Fig. S1B**). We therefore reasoned that acarbose likely differentially inhibits a Sus enzyme between the two organisms. To ascertain if acarbose targets the Sus machinery to elicit its inhibitory effects on Bt, and to a lesser extent on Bo, BoΔ*bovatus_03809-03803* and BtΔ*susABCDEFG*, hereafter referred to as BoΔSus and BtΔSus, respectively, were grown in the presence or absence of 50 μM acarbose. Note that both BoΔSus and BtΔSus cannot grow in minimal media with maltoheptaose or starch but do grow on maltose (**Fig. 2C**, **Fig. S1C**) (39). Although ΔSus strains lag on maltose compared to the respective wild-type strain, the addition of acarbose did not extend this lag or change growth kinetics (**Fig. 2C,D**). Likewise, no additional lag was observed for the ΔSus strains grown in glucose and acarbose (Fig. S1D). These data led us to hypothesize that a difference in the Bo and Bt Sus was responsible for Bo’s resistance to acarbose induced growth inhibition.

### Sus outer membrane α-amylases are not the source of the different Bo and Bt acarbose phenotypes

The most likely candidate for differential inhibition is the extracellular α-amylase enzymes, as this is the most obviously different component of the loci (**Fig. 1C**). BoGH13A_Sus_ and SusG are both α-amylases belonging to the glycoside hydrolase family 13 (GH13). BoGH13A_Sus_ belongs to subfamily 47 whereas SusG belongs to subfamily 36. Furthermore, they have vastly different structural properties (38, 51) (**Fig. 3A,B**). BoGH13A_Sus_ has two N-terminal carbohydrate binding modules (CBMs): CBM98, which binds maltooligosaccharides, and CBM48, which does not (**Fig. 3A**) (51). SusG features a CBM58 that interrupts its catalytic domain (**Fig. 3B**) (38). Both enzymes have A, B, and C domains that are GH13 hallmarks although the BoGH13A_Sus_ B domain is much smaller than SusG’s (53). BoGH13A_Sus_ and SusG have signal peptides that destine them to the outer leaflet of the outer membrane via lipidation at an N-terminal cysteine residue. The orientation of the SusG GH13 domain, however, is opposite that of BoGH13A_Sus_ relative to the outer membrane (**Fig. 3A,B**). Given the extensive differences between BoGH13A_Sus_ and SusG at a structural level, we hypothesized that SusG would be more inhibited by acarbose than BoGH13A_Sus_ in congruence with the respective organism’s acarbose growth phenotype.

**Figure 3.**
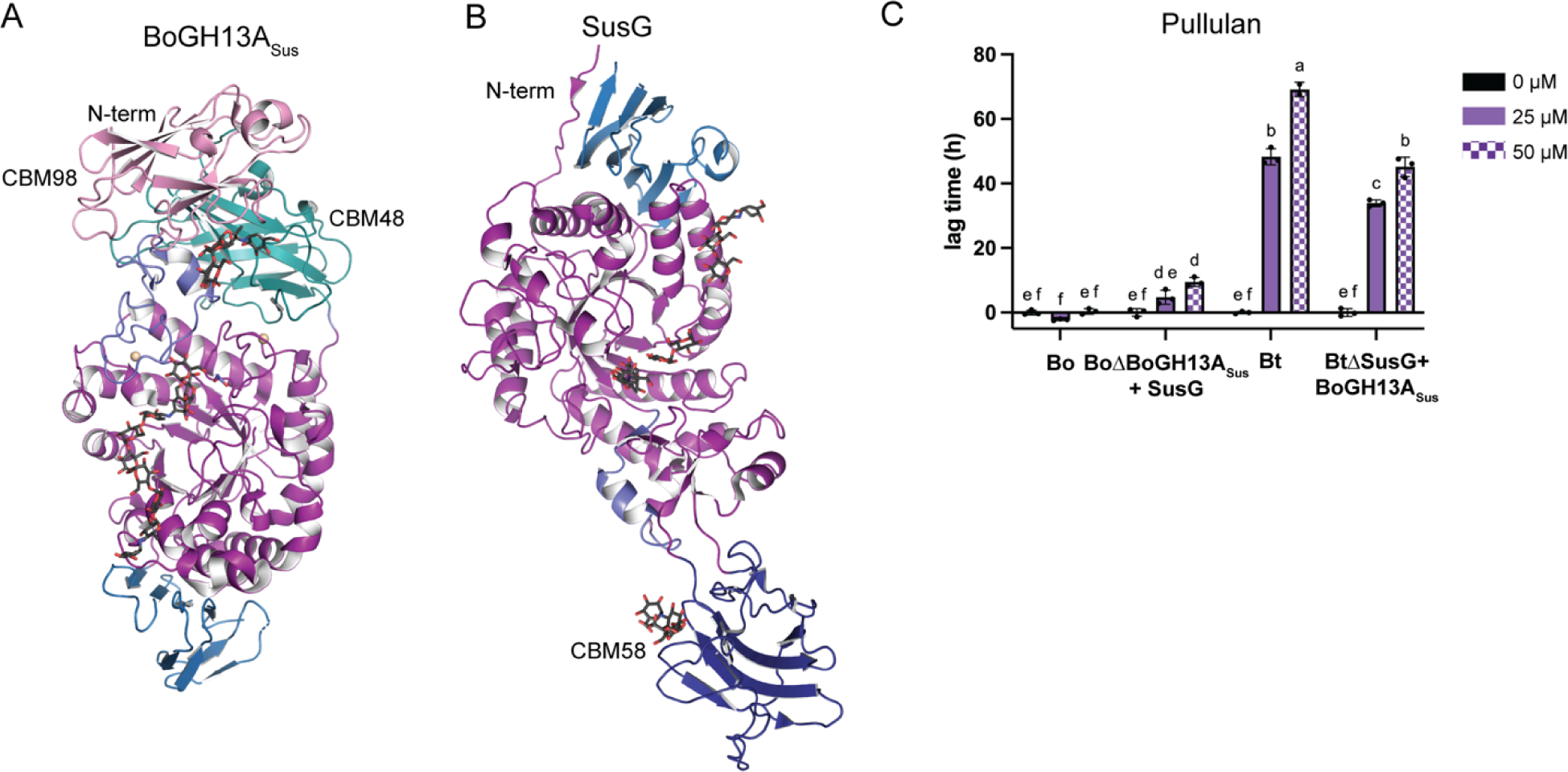
The extracellular amylases BoGH13A_Sus_ and SusG are not the source of the different acarbose phenotypes in Bo and Bt, respectively. A) The crystal structure of BoGH13A_Sus_ bound to acarbose (PDB: 8DL2 (51)). The GH13 A domain is in purple, B domain in slate, and C domain in blue. CBM98 is pink and CBM48 is teal. B) The crystal structure of SusG bound to acarbose (PDB: 3K8M (38)). The GH13 domains are colored as in A. CBM58 is dark blue. They are oriented with their N-termini at the top of the figure, as both are anchored to the outer membrane via lipidation of an N-terminal Cys. Figures were rendered in PyMOL (54). C) Given that BoGH13A_Sus_ does not optimally complement BtΔSusG, these growths were performed with bacteria pre-grown on minimal media (MM) + 5 mg/ml maltose to induce *sus* expression, then inoculated into MM + 2.5 mg/ml pullulan in the indicated acarbose concentrations. Acarbose induced lag times are shown. Statistical analyses were performed using a two-way ANOVA using a cutoff of p≤0.05. Conditions with the same letter(s) were not significantly different from one another.

To quantify acarbose inhibition kinetics of each enzyme, IC_50_ values were determined using a fluorescent starch assay. To our surprise, BoGH13A_Sus_ was more inhibited by acarbose than SusG, borne out by a 10-fold lower IC_50_ (**Table 2**, **Fig. S2A,B**). The SusG IC_50_ is in rough alignment with the growth derived IC_50_s for polysaccharides in **Table 1**, but the IC_50_ for BoGH13A_Sus_ is much lower than acarbose concentrations tolerated by Bo during growth. In previous work we showed that SusG displays a minimal amount of acarbose hydrolysis *in vitro* over time using elevated enzyme concentrations, but BoSusG cannot break down acarbose (38, 51).

**Table 2.** Enzyme Kinetic Parameters.

| Protein | $K_M$ (mM) | $k_{cat}$ (s<sup>-1</sup>) | $k_{cat}/K_M$ (s<sup>-1</sup>mM<sup>-1</sup>) | Inhibition Constant |
| --- | --- | --- | --- | --- |
| BoGH13A<sub>Sus</sub> | N/A | N/A | N/A | 2.2 ± 0.35 μM<sup>a</sup> |
| SusG | N/A | N/A | N/A | 68.2 ± 13.8 μM<sup>a</sup> |
| BoGH13B<sub>Sus</sub> | 0.135 ± 0.010 | 18.2 ± 0.5 | 135 | 122.6 ± 9.2 nM<sup>b</sup> |
| SusA | 0.117 ± 0.009 | 13.8 ± 0.3 | 120 | 94.7 ± 6.9 nM<sup>b</sup> |
| BoGH97C<sub>Sus</sub> | 0.078 ± 0.015 | 21.5 ± 1.2 | 276 | 69.3 ± 8.9 nM<sup>c</sup> |
| SusB | 0.089 ± 0.013 | 23.9 ± 1.1 | 269 | 53.5 ± 12.5 nM<sup>c</sup> |
| BoGH97D | 0.088 ± 0.015 | 51.2 ± 2.8 | 582 | 133.8 ± 22.4 nM<sup>c</sup> |
| BtGH97H | 0.088 ± 0.014 | 48.6 ± 2.5 | 552 | 162.4 ± 24.3 nM<sup>c</sup> |

<sup>a</sup>IC<sub>50</sub>
<sup>b</sup>αK<sub>i</sub>
<sup>c</sup>K<sub>i</sub>

To substantiate our *in vitro* findings, the extracellular α-amylase genes were swapped between Bo and Bt in their respective *sus* loci to test whether the acarbose phenotype tracked with the expressed enzyme. BoΔBoGH13A_Sus_ and BtΔSusG cannot grow on starch polysaccharides (32, 36, 51). We have previously demonstrated that *susG* complements a BoΔBoGH13A_Sus_ strain on potato starch, potato amylopectin, pullulan, and glycogen (51). However, *bovatus_03803* (the gene for BoGH13A_Sus_) poorly complements BtΔ*susG* growth on potato amylopectin and glycogen but restores growth on pullulan and to a lesser extent, potato starch, when cells are pre-grown on maltose to induce *sus* expression (51). Therefore, acarbose induced growth lags on pullulan in the BoΔBoGH13A_Sus_+SusG and BtΔSusG+BoGH13A_Sus_ strains were assessed. Both strains grew normally in glucose (51) (**Fig. S2C**). BoGH13A_Sus_ imparted a small benefit to BtΔSusG compared to Bt and BoΔBoGH13A_Sus_+SusG lagged slightly more compared to Bo, both with statistical significance. However, the swapped strains nonetheless responded to acarbose more like the parent strain and without changing the overall phenotype (**Fig. 3C**). Together with the inhibition data, this suggests that the different Bo and Bt responses to acarbose are not primarily driven by BoGH13A_Sus_ or SusG.

### Acarbose likely competes with maltooligosaccharides for transport through BoSusC and SusC

BoΔBoGH13A_Sus_ and BtΔSusG can grow on short maltooligosaccharides (32, 36, 37, 51). Therefore, to examine acarbose induced growth inhibition in the absence of BoGH13A_Sus_ and SusG, we grew the deletion mutants on various maltooligosaccharides with or without acarbose. In the absence of BoGH13A_Sus_ or SusG, acarbose inhibited growth in a size-dependent manner with growth on shorter maltooligosaccharides displaying a longer lag time (**Fig. 4A**). Acarbose did not affect growth on glucose (**Fig. S3A**). Furthermore, the shape of the oligosaccharide influenced growth. Both Bo and Bt grew on α-cyclodextrin (αCD) independent of BoGH13A_Sus_ and SusG, respectively. However, WT and ΔGH13_E_ (deletion of the extracellular Sus GH13s) strains of Bo and Bt did not grow on αCD in the presence of acarbose (**Fig. 4B**). Taken together, these results suggest that 1) acarbose may compete with maltooligosaccharides for transport through BoSusC and SusC in a size and shape dependent manner (**Fig. 4C**) and 2) acarbose can enter the periplasm to inhibit Sus periplasmic enzymes.

**Figure 4.**
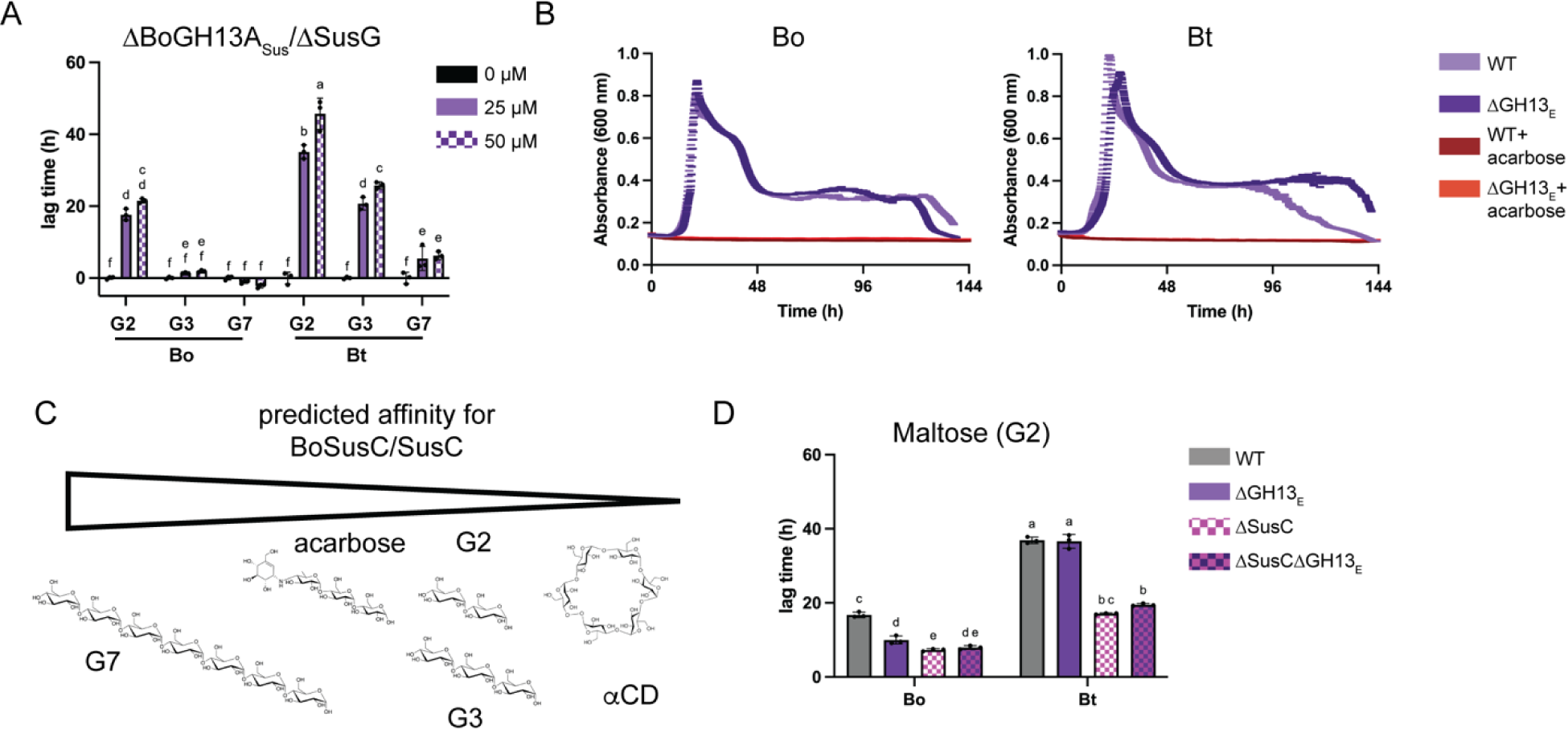
Acarbose likely competes with maltooligosaccharides for transport through BoSusC and SusC. A) Bacteria were pre-grown in minimal media (MM) with glucose and back diluted in MM + 2.5 mg/l maltose (G2), maltotriose (G3), or maltoheptaose (G7) with and without the indicated acarbose concentrations. Acarbose induced lag times are shown. B) Bacteria were grown as described in Fig. 2B in 2.5 mg/ml α-cyclodextrin +/− 50 µM acarbose. ΔGH13_E_ (GH13 extracellular) corresponds to BoΔBoGH13A_Sus_ and BtΔSusG. C) Based on A and B, we predict that maltoheptaose has the highest affinity for BoSusC/SusC whereas αCD has the lowest. D) Bacteria were pre-grown in MM and back diluted in MM + 2.5 mg/ml maltose +/− 50 µM acarbose. ΔGH13_E_ (GH13 extracellular) corresponds to BoΔBoGH13A_Sus_ and BtΔSusG. Acarbose induced lag times are shown. Statistical analyses in A and D were performed with a two-way ANOVA. Conditions with the same letter(s) were not significantly different from one another. A cutoff of p≤0.05 was used.

To probe the first possibility, strains lacking BoSusC/SusC were grown in maltose with or without acarbose since growth on maltose, but not longer maltooligosaccharides, does not require SusC in Bt (**Fig. S3B**) (36, 39). In the mutants lacking this transporter, cells lagged significantly less in the presence of maltose plus acarbose compared to the WT cells (**Fig. 4D**). We hypothesize that this reduced lag is because maltose transport is BoSusC/SusC independent whereas acarbose transport, due to its larger size, is SusC/BoSusC dependent. ΔBoSusC and ΔSusC strains exhibited a growth benefit in glucose with acarbose compared to the WT strains (**Fig. S3C**). Even though Bo and Bt aren’t affected by acarbose when grown in glucose, there may be some benefit to keeping acarbose outside of the cell in the strains lacking an outer membrane Sus transporter. Finally, discrepant Bo and Bt acarbose phenotypes were still observed in the absence of the BoSusC/SusC transporter and extracellular amylase (**Fig. 4D**). Together these data supported that the target of acarbose inhibition is intracellular, as deletion of the BoSusC/SusC transporters provided partial relief of inhibition, and the surface amylases do not drive the difference in phenotype between Bo and Bt.

### Periplasmic GH13 enzymes are not the source of the different Bo and Bt acarbose phenotypes

Because deletion of the *sus* locus relieves the acarbose-induced growth lag in MM plus maltose, we next examined the influence of the periplasmic Sus enzymes in mediating sensitivity to the drug. The genes encoding BoGH97C_Sus_ and SusB were mutated to introduce a stop codon after eight amino acids, as an in-frame deletion of *susB* has polar effects in Bt (40). While *susA* is transcribed separately from *susBCDEFG* (40), genes for BoGH13B_Sus_ and SusA were also knocked out via an early stop codon for consistency. Deleting BoGH13B_Sus_ from Bo and SusA from Bt did not rescue the acarbose induced lag phenotype when cells were grown on maltose and neither strain lagged in glucose (**Fig 5A**, **Fig. S4A**). BtΔSusA grew in maltoheptaose and amylopectin but still exhibited an acarbose induced growth lag (40, 55) (**Fig. 5B,C**). Conversely, Bo demonstrated a very small growth lag in 50 μM acarbose when grown on maltoheptaose or amylopectin. Deleting BoGH13B_Sus_ rendered the organism slightly more susceptible to acarbose, although the difference was not statistically significant, rather than relieving the already miniscule growth lag (**Fig. 5B,C**).

**Figure 5.**
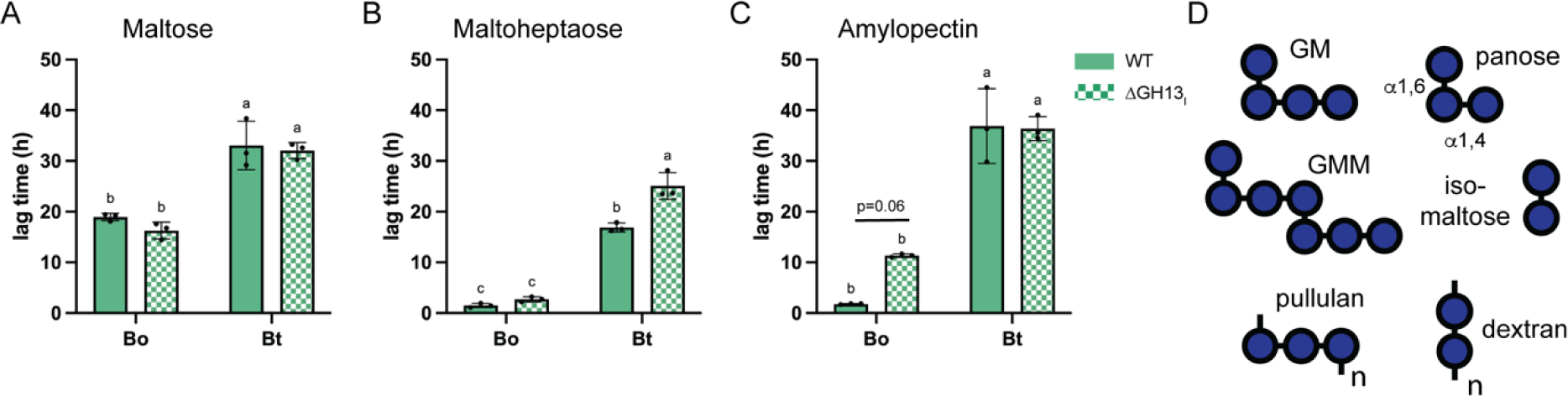
Periplasmic Sus GH13 enzymes do not underpin the Bo and Bt responses to acarbose. A-C) Bo and Bt were pre-grown in minimal media (MM) with glucose and back diluted into MM + 2.5 mg/ml maltose, maltoheptaose, or amylopectin +/− 50 µM acarbose. ΔGH13_I_ (GH13 intracellular) corresponds to BoΔBoGH13B_Sus_ and BtΔSusA. Acarbose induced growth lags are displayed and statistical analyses were performed with a two-way ANOVA. Conditions with the same letter(s) were not significantly different from one another. A cutoff of p≤0.05 was used. D) Cartoon renderings of the α-glucans used for TLC experiments with the periplasmic GH13 and GH97 enzymes.

To corroborate that BoGH13B_Sus_ and SusA are not the primary drivers of the different acarbose responses, the enzymes were recombinantly expressed in *E. coli* and purified to compare their activities. SusA is known to target α1,4 linkages in maltose through maltoheptaose (G2 – G7), β-cyclodextrin (βCD), and pullulan (producing panose as a product) and does not recognize α1,6 bonds (41) (**Fig. 5D**). Substrate preferences of BoGH13B_Sus_ were qualitatively compared to those in SusA by overnight incubation with various substrates and analyzed via thin layer chromatography. Both proteins exclusively targeted the α1,4 linkages in starch oligosaccharides as panose, isomaltose, and dextran (polymer of α1,6 glucans) were not broken down (**Fig. 5D**, **S4B,C**). To break down linear α1,4 glucans, a minimum of 3 active site subsites must be filled as neither enzyme broke down maltose. Both enzymes released maltose and glucose as major products, consistent with their function as endo-acting enzymes (41). BoGH13B_Sus_ and SusA are annotated as neopullulanases with cyclomaltodextrinase N- and C-terminal domains and belong to the recently identified GH13_46 subfamily (56).

Both enzymes broke down pullulan (into panose), αCD, βCD, the pullulan oligosaccharides 6^3^-α-d-glucosyl-maltotriose (GM), 6^3^-α-d-glucosyl-maltotriosyl-maltotriose (GMM), and amylopectin (**Fig. 5D**, **S4B,C**). Neither was active on glycogen and BoGH13B_Sus_ was more active than SusA on potato starch. That either enzyme was active on polysaccharide is somewhat surprising since these are periplasmic enzymes. Our previous work demonstrated that SusC preferentially transports larger maltooligosaccharides of 15-17 Glc units, and this relative size was corroborated in structural studies of the levan SusCD-like transporter (37, 57). Thus, BoGH13B_Sus_ and SusA likely encounter relatively long maltooligosaccharides in the periplasm. Most importantly, BoGH13B_Sus_ and SusA did not break down acarbose (**Fig. S4B,C**).

BoGH13B_Sus_ and SusA inhibition by acarbose was quantified using pNP-maltohexaose (pNP-G6) as a substrate, for which the enzymes have comparable catalytic efficiency (**Table 1**, **Fig S5A,B**). To our surprise, acarbose modelled best as an uncompetitive inhibitor of both enzymes, implying that acarbose only binds to the enzyme-substrate complexes and not to free enzyme (**Fig S5C,D**). This was unexpected since acarbose is typically a mixed or a competitive enzyme inhibitor (58, 59). Furthermore, in cases where acarbose uncompetitively inhibits amylase activity, it does so during polysaccharide, not oligosaccharide, breakdown (59). However, even though our kinetic data was best modeled via uncompetitive inhibition, we are inclined to conclude that acarbose is instead a mixed inhibitor of these enzymes because it binds to free enzyme as measured by isothermal titration calorimetry (**Fig. S6A-D**). Wild-type enzyme has two acarbose binding sites whereas a catalytically inert variant of each enzyme (D331N) has one (**Table S1**). It is tempting to speculate that active site binding opens a second acarbose binding site, but this site could not be predicted based on the structures of similar enzymes in the Protein Databank (PDB). Both BoGH13B_Sus_ and SusA are dimers (**Fig. S6E**), raising the possibility of allosteric modulation. Nonetheless, the inhibition constants are the same order of magnitude (α*K*_i_s of 123 vs 95 nM for Bo and Bt enzymes, respectively, **Table 2**), suggesting that BoGH13B_Sus_ and SusA do not explain the Bo and Bt responses to acarbose.

### Periplasmic Sus GH97s drive acarbose responses but do not underpin the different Bo and Bt phenotypes

Deletion of the genes encoding the periplasmic Sus GH97s (BoΔBoGH97C_Sus_ and BtΔSusB) eliminated growth on amylopectin (**Fig. S7A,B**). Our results with Bt differed on this point from previous work by the Salyers lab but are in accordance with data collected from barcoded transposon libraries of Bt (40, 55, 60). Without acarbose, BtΔSusB exhibited a significant growth lag compared to Bt on maltose, maltotriose, and maltoheptaose while BoΔBoGH97C_Sus_ lagged on maltoheptaose compared to Bo but not maltose or maltotriose (**Fig. 6A**). BtΔSusB and BoΔBoGH97C_Sus_ were nonetheless screened for acarbose-induced growth inhibition in all three oligosaccharides. Neither deletion strain exhibited an acarbose induced growth lag when grown in glucose (**Fig. S7C**). In the absence of BoGH97C_Sus_ and SusB, Bo and Bt no longer had an acarbose induced lag in maltose (**Fig. 6B**). Compared to WT, the acarbose induced growth lag in maltotriose was rescued in BtΔSusB but BoΔBoGH97C_Sus_ grew similarly to WT Bo (**Fig. 6C**). Strikingly, when grown on maltoheptaose, BtΔSusB did not have a growth defect, but BoΔBoGH97C_Sus_ growth was impaired by acarbose (**Fig. 6D**). Nonetheless, we swapped the GH97 genes between organisms in the context of their *sus* loci in hopes of swapping acarbose phenotypes. Instead, BtΔSusB+BoGH97C_Sus_ and BoΔBoGH97C_Sus_+SusB grow like their wild-type counterparts with or without the outer membrane GH13s which are active on maltoheptaose (BoGH13A_Sus_) and maltoheptaose + maltotriose (SusG) (**Fig. 6B-D**) (38, 51). BoΔBoGH97C_Sus_+SusB and BoΔBoGH13A_Sus_ΔBoGH97C_Sus_+SusB had a slightly smaller lag time compared to WT Bo when grown in maltose, but not maltotriose or maltoheptaose (**Fig. 6B-D**).

**Figure 6.**
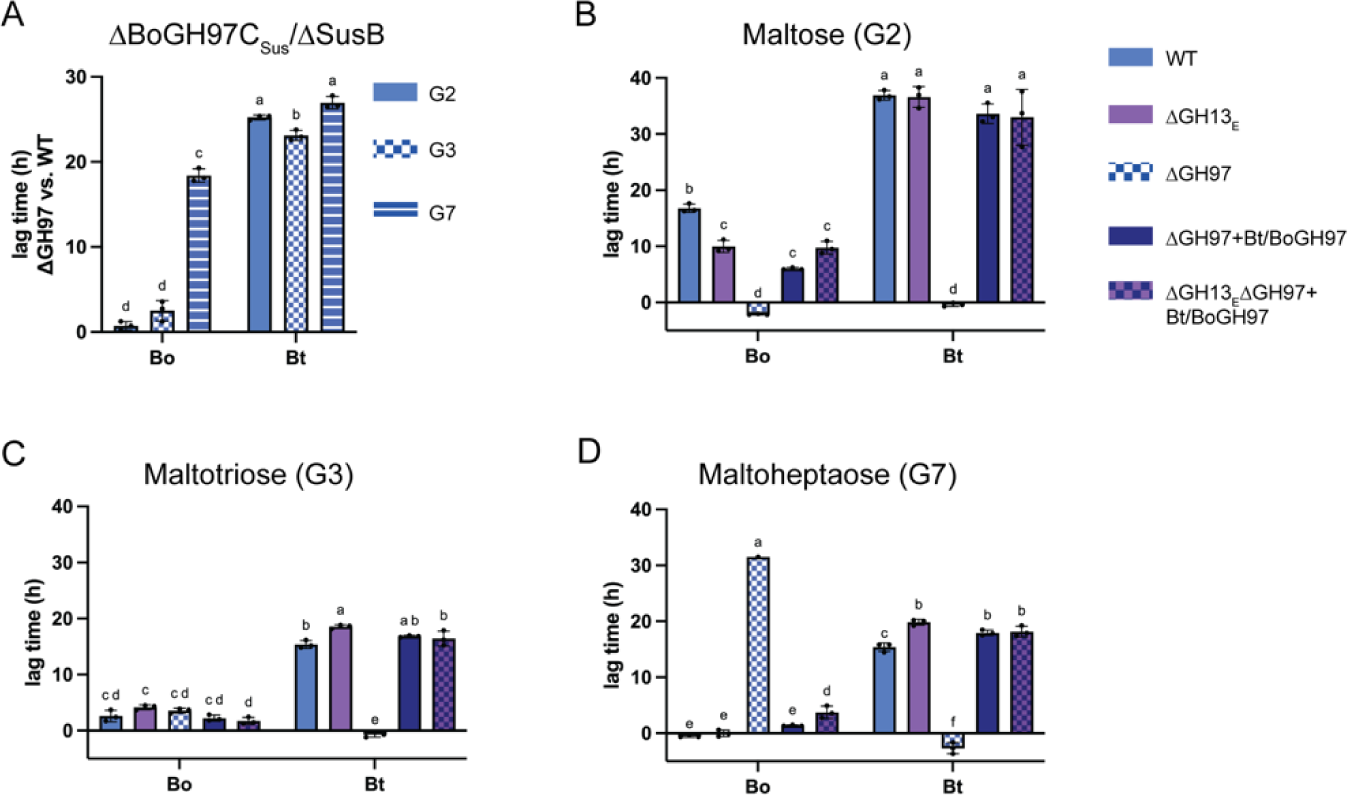
Periplasmic Sus GH97 enzymes are the primary acarbose target but do not explain the different Bo and Bt phenotypes in acarbose. A) Bacteria were grown in minimal media (MM) with glucose and back diluted into MM + 2.5 mg/ml maltose, maltotriose, or maltoheptaose. Difference in time to OD_600_ of 0.3 between WT Bo and BoΔGH97C_Sus_ and between WT Bt and BtΔSusB is shown. B-D) Bacteria were pre-grown in MM with glucose and back diluted into MM + 2.5 mg/ml maltose, maltotriose, or maltoheptaose +/− 50 µM acarbose. ΔGH13_E_ (GH13 extracellular) corresponds to BoΔBoGH13A_Sus_ and BtΔSusG. ΔGH97 corresponds to BoΔBoGH97C_Sus_ and BtΔSusB. Acarbose induced growth lags are displayed and statistical analyses were performed with a two-way ANOVA. Conditions with the same letter(s) were not significantly different from one another. A cutoff of p≤0.05 was used.

Furthermore, BoGH97C_Sus_ and SusB appear to have similar substrate preferences. Our work here, and previously published work on SusB, demonstrate that both enzymes are exo-acting glucoamylases that accommodate α1,6 and α1,4 linkages, and do not break down acarbose (41, 42) (**Fig. S7D**). During growth on α-glucans, BoGH97C_Sus_ and SusB are ultimately responsible for degrading oligosaccharides not recognized by BoGH13B_Sus_ and SusA (maltose, isomaltose, panose) and vice versa. BoGH97C_Sus_ and SusB cannot break down cyclodextrins, but BoGH13B_Sus_ and SusA have this activity (**Fig. S7D**).

The kinetic parameters of SusB and BoGH97C_Sus_ using pNP-glucose as a substrate demonstrated they have similar catalytic efficiencies and the SusB *K*_i_ (54 nM) for acarbose was comparable to published work (150 nM and 108) (**Table 2**, **Fig. S8A-D**) (42, 43). Remarkably, BoGH97C_Sus_ had a similar *K*_i_ (69 nM). SusB has been structurally characterized and is known to be competitively inhibited by acarbose (42, 43). We solved a co-crystal structure of BoGH97C_Sus_ with acarbose bound in the active site near the dimer interface (**Table S2**). Acarbose binds identically to the SusB and BoGH97C_Sus_ active sites (**Fig. S8E**). Collectively, these data suggest that although BoGH97C_Sus_ and SusB drive most of the acarbose phenotype in Bo and Bt, respectively, they are not responsible for the acarbose induced phenotypic differences.

### Bo expresses a GH97 enzyme whose homolog is not expressed by Bt

Because Bo and Bt ΔSus strains no longer exhibit a growth lag due to acarbose when grown in maltose (**Fig. 2C**), we had hypothesized that the differences in Sus were responsible for the phenotypic differences between Bo and Bt during growth on acarbose. However, the enzymes encoded in the *sus* of both organisms have nearly identical responses to acarbose and swapping these components does not convert the phenotypes. Thus, we concluded that while SusB is a significant target of acarbose for both organisms, we hypothesized that Bo has an additional mechanism to subvert acarbose growth inhibition for two reasons. First, we noticed that in the absence of acarbose, BoΔSus consistently grows up faster than BtΔSus in maltose, suggesting it might harbor a glucosidase/glucoamylase not found in Bt (**Fig. 2D**). Second, also in the absence of acarbose, BoΔBoGH97C_Sus_ did not exhibit a growth lag on maltose or maltotriose compared to WT Bo but BtΔSusB did (**Fig. 6A**).

To test our hypothesis, WT and ΔSus strains of Bo and Bt were grown in maltose and harvested at similar ODs. Bacteria were pelleted, sonicated, and centrifuged to clarify debris. The clarified lysates were assayed for pNP-glucose activity using 1 mM substrate. As expected, wild-type lysates from both organisms exhibited robust pNP-glucose activity due to BoGH97C_Sus_ and SusB (**Fig. 7A**). BtΔSus recapitulated published work in a BtΔSusB strain and had minimal pNP-glucose activity (40). BoΔSus, however, had activity comparable to WT Bo, suggesting it harbors one or more non-Sus backup glucosidases/glucoamylases (**Fig. 7A**). To our surprise, this activity was inhibited by 1 µM acarbose in the conditions tested (**Fig. 7A**).

**Figure 7.**
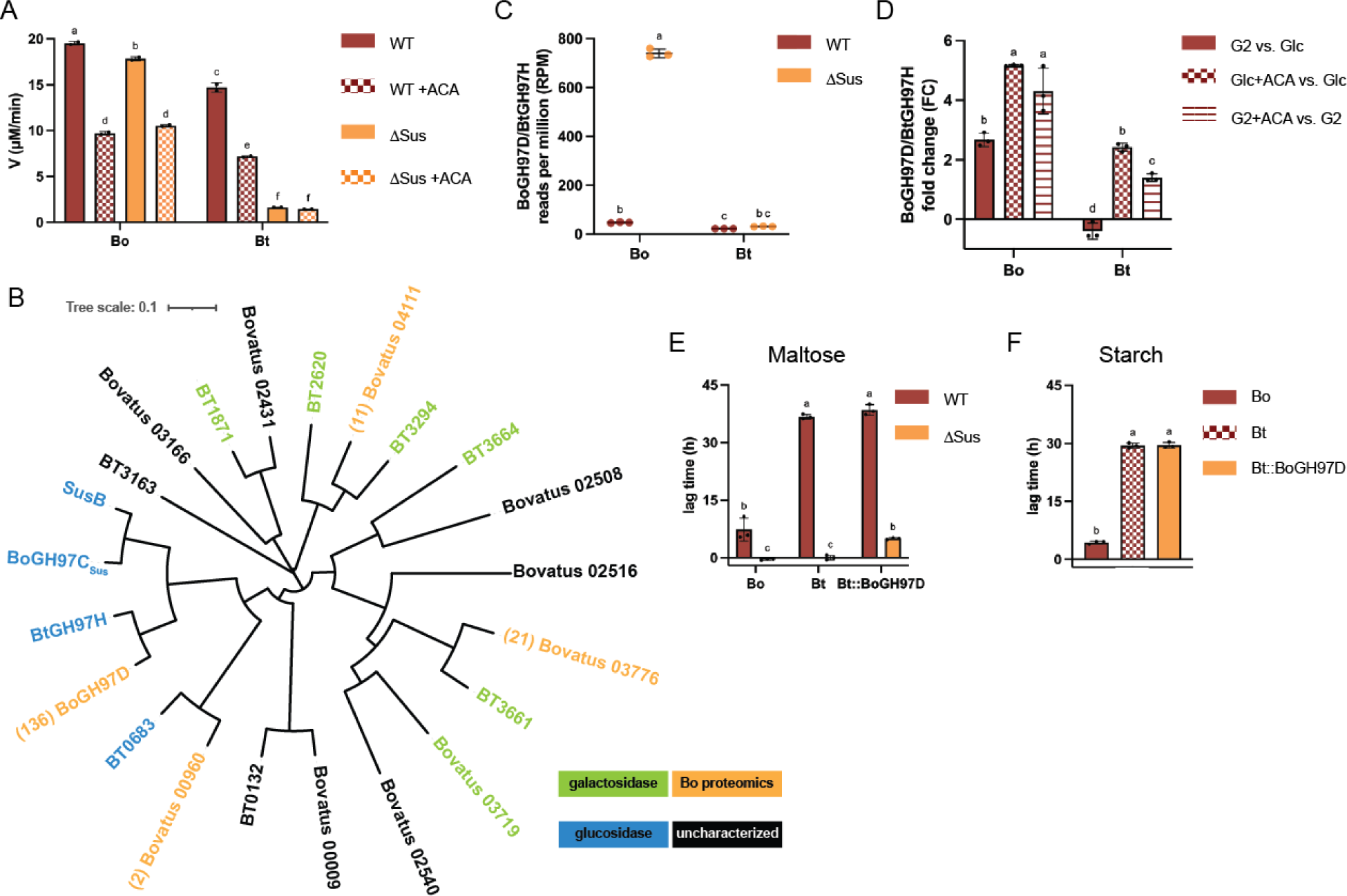
Bo upregulates a non-Sus GH97, BoGH97D, that does not rescue the acarbose induced growth lag when expressed by Bt. A) WT and ΔSus strains of Bo and Bt were grown in minimal media (MM) + 5 mg/ml maltose to the same OD_600_ and pelleted. Pellets were washed in PBS and cells were sonicated to release contents. Lysates were assayed in 1 mM pNP-Glc +/− 1 µM acarbose. B) BoΔSus cellular contents following growth in maltose were purified according to the Methods to increase specific activity on pNP-Glc. Proteins in the purest and most active fractions were identified via LC-MS/MS. The most abundant predicted α-glucan active enzyme was bovatus_04772 (BoGH97D). A phylogenetic tree of all Bo and Bt GH97s is shown along with the number of peptide to spectrum (PSM) matches observed in the proteomics experiment. Locus tags with colored names have previously been characterized biochemically (40, 42, 43, 63–68) or were characterized for this study (BoGH97D, BtGH97H). C) WT and ΔSus strains of Bt and Bo were grown in MM + 5 mg/ml maltose. Total RNA was purified and subjected to RNAseq. Reads per million are displayed. D) Bo and Bt were grown in MM with 5 mg/ml glucose or maltose +/− 50 µM acarbose and total RNA was purified. qPCR was performed to determine the fold change in BoGH7D and BtGH97H expression in the following conditions: maltose (G2) vs. glucose (Glc); glucose + 50 µM acarbose vs. glucose; maltose + 50 µM acarbose vs. maltose. E,F) The indicated strains were pre-grown on MM + 5 mg/ml maltose to recapitulate growths used for RNAseq and otherwise grown in MM + 2.5 mg/ml maltose or starch +/− 50 µM acarbose. All statistical analyses were performed with a two-way ANOVA. Conditions with the same letter(s) were not significantly different from one another. A cutoff of p≤0.05 was used.

We used two complementary approaches to identify Bo glucosidases. First, activity guided fractionation was performed from BoΔSus grown in maltose. Following sonication and centrifugation, clarified lysates were subjected to ammonium sulfate cuts and chromatographic separation. Throughout, fractions were tested for pNP-glucose activity and the identities of proteins in the purest and most active fractions were determined via LC MS/MS (**Table S3**). The most abundant enzyme that we predicted would exhibit pNP-glucose activity was Bovatus_04772 (hereafter referred to as BoGH97D), annotated as a GH97. Of other glycoside hydrolase families with α-glucan activity (GH13, GH15, GH31, GH57, and GH77), two GH31s were identified by a single peptide for each (61). Bo does not encode any other predicted α-glucan active GHs such as maltose phosphorylases (GH65), GH4, GH63, or GH122s in its genome (61). Furthermore, of the four GH97s identified by proteomics, two are predicted α-galactosidases and the other predicted α-glucosidase had a single peptide sequenced (**Fig. 7B**, **Table S3**). Bt, however, encodes a homolog of BoGH97D, Bt4581 (hereafter referred to as BtGH97H), which is 90% identical.

The other unbiased approach we took to identify the Bo non-Sus enzyme with pNP-Glc activity was RNAseq of WT and ΔSus strains of Bo and Bt grown in maltose. The BoGH97D transcript was nearly 16-fold more abundant in BoΔSus compared to WT Bo whereas the BtGH97H transcript was only 1.5-fold more abundant in BtΔSus compared to WT Bt (**Fig. 7C**, **Table S4**). Furthermore, no Bt transcripts for GH31s or GH97s passed our significance or fold-change threshold (p ≤ 0.05; 2-fold change, **Table S5**). While Bovatus_02431, a GH97 with predicted α-galactosidase activity based on phylogeny, did exceed our threshold, there was one tenth as much transcript as BoGH97D, and was not identified in the proteomics experiments (**Fig. 7B**, **Tables S3** and **S4**). A single Bo GH31, Bovatus_03177, also passed our cutoff but is forty times less abundant than BoGH97D transcript and was not identified by proteomics (**Tables S3** and **S4**).

BoGH97D and BtGH97H were purified and confirmed to be α-glucosidases and not α-galactosidases (**Fig. S9A**). They break down all α1,4 and α1,6 linked substrates that BoGH97C_Sus_ and SusB are active on (**Fig. S9B**) BoGH97D and BtGH97H are predicted to localize to the periplasm due to their SPI signal peptides (62). We did not confirm whether they are retaining or inverting enzymes (and thus should be considered glucosidases or glucoamylases according to (42)). Based on comparisons to known determinants of inverting activity in SusB, we predict BoGH97C_Sus_, BoGH97D, and BtGH97H to be inverting (**Fig. S9C**) (43). BoGH97D and BtGH97H had comparable catalytic efficiencies using pNP-Glc as a substrate, and both enzymes are competitively inhibited by acarbose with comparable *K*_i_s (134 vs 162 nM for BoGH97D and BtGH97H, respectively) (**Table 2**, **Fig. S10A-D**). Since both BoGH97D and BtGH97H are active enzymes, we reasoned that if BoGH97D is at least partly responsible for the acarbose phenotype in Bo, it is regulated differently than BtGH97H. BoGH97D was upregulated by maltose and acarbose compared to glucose alone (**Fig. 7D**). Conversely, BtGH97H was not upregulated by maltose but was upregulated by acarbose compared to glucose alone. However, acarbose upregulated BoGH97D over two times as much as BtGH97H when bacteria were grown in glucose. When grown in maltose, BtGH97H was not upregulated by the addition of acarbose, but BoGH97D was (**Fig. 7D**). These data suggest that BoGH97D is upregulated by acarbose in the context of growth on starch but that BtGH97H is not.

### BoGH97D does not confer acarbose resistance to Bt

Our attempts to delete BoGH97D from the Bo genome were unsuccessful so we could not assess its contribution to acarbose resistance in Bo. Instead, we overexpressed BoGH97D constitutively in Bt and BtΔSus to see if it rescued part of the acarbose induced lag phenotype (69). First, we repeated the previously described lysate experiments and showed that BtΔSus::BoGH97D lysates had more pNP-Glc activity than BtΔSus ones, but not as much as BoΔSus lysates (**Fig. S11A**). Bt::BoGH97D lysates did not have more glucosidase activity than Bt lysates, most likely due to the amount of SusB in the WT lysates. Nonetheless, we screened these Bt strains for resistance to acarbose. Bt::BoGH97D and BtΔSus::BoGH97D behaved similarly to Bt and BtΔSus in the absence of acarbose when grown in maltose or starch (**Fig. S11B,C**). Neither strain exhibited an acarbose induced growth lag in glucose (**Fig. S11D**). BtΔSus::BoGH97D grew similarly to BtΔSus in maltose and Bt::BoGH97D grew similarly to Bt in maltose and starch (**Fig. 7E,F**). Thus, BoGH97D did not provide Bt any growth advantage in the presence of acarbose, suggesting that differential regulation of the non-Sus enzymes BoGH97D and BtGH97H does not contribute significantly to the different acarbose phenotypes between Bo and Bt.

### Acarbose and maltose upregulate BoSus to a greater extent than BtSus

We have shown that acarbose inhibits all Sus enzymes as well as the non-Sus glucosidases BoGH97D and BtGH97H. Acarbose also likely competes with maltooligosaccharides for transport through BoSusC/SusC. An additional periplasmic component we have not addressed is the regulator BoSusR/SusR. We wondered if Bo and Bt upregulate their *sus* loci differently in response to maltose and acarbose. To test this, qPCR on the genes for BoSusC and SusC was performed. To our surprise, compared to glucose alone, acarbose and glucose upregulated *sus* more than maltose in both organisms. 50 µM acarbose was used in these experiments whereas the bacteria were grown in 5 mg/ml maltose (∼14.6 mM). Furthermore, the magnitude of responses between Bo and Bt was strikingly different as maltose upregulated BoSus ∼210-fold and BtSus ∼35-fold. The addition of 50 µM acarbose to the glucose culture upregulated BoSus ∼335-fold and BtSus ∼80-fold. On the other hand, the addition of 50 µM acarbose to cultures grown in maltose only upregulated Bo and BtSus ∼5-fold compared to maltose alone (**Fig. 8A**).

**Figure 8.**
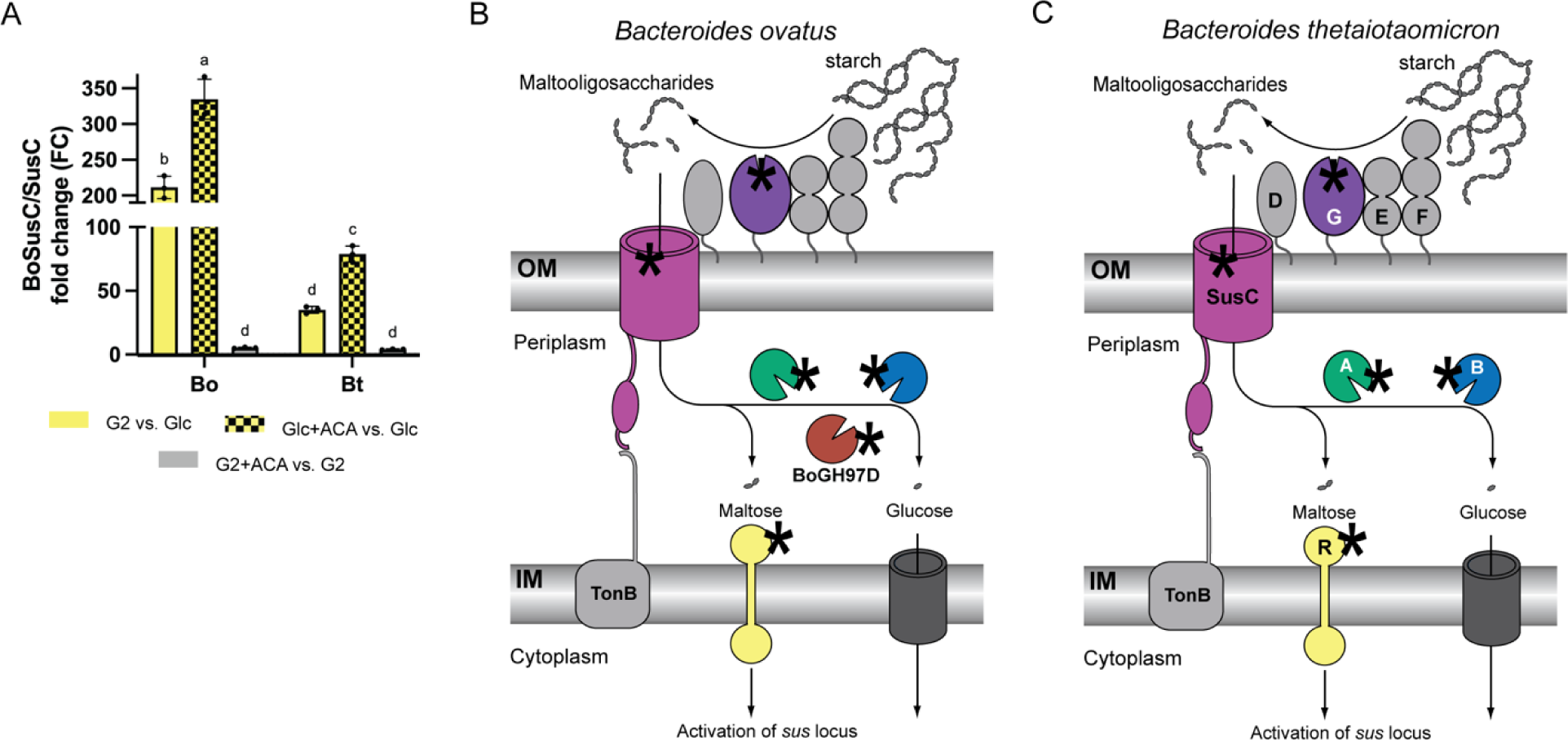
Acarbose targets BoSus and BtSus components from the outside of the cell to the periplasmic *sus* regulator. A) Bo and Bt were grown in 5 mg/ml glucose or maltose +/− 50 µM acarbose and total RNA was purified. qPCR was performed to determine the fold change in BoSusC and SusC expression in the following conditions: maltose (G2) vs. glucose (Glc); glucose + 50 µM acarbose vs. glucose; maltose + 50 µM acarbose vs. maltose. Statistical analyses were performed with a two-way ANOVA. Conditions with the same letter(s) were not significantly different from one another. A cutoff of p≤0.05 was used. B) Model for how acarbose interacts with BoSus, including the addition of BoGH97D. C) Model for how acarbose interacts with BtSus. Asterisks in B and C represent acarbose.

## Discussion

Acarbose is an effective type 2 diabetes medication that is showing promise for potential anti-aging and cardioprotective effects. Acarbose exposure also affects the gut microbiome. Among these effects is a reduction in the relative abundance of *Bacteroides* species, including *Bacteroides thetaiotaomicron* (Bt), *Phocaeicola dorei*, and *Phocaeicola vulgatus* (10). Acarbose impairs the growth of all three species when using starch *in vitro* as a carbon source (24, 25) (**Fig. 1D**). Here, we fill in some of the mechanistic gaps between exposure to acarbose and changes in the gut microbiome and host. Given that *Bacteroides ovatus* (Bo) and Bt exhibit very different acarbose induced growth lag phenotypes (**Fig. 2B**, **Fig. S1A**), we initially hypothesized that a Sus enzyme was differentially inhibited. However, the periplasmic Bo and Bt Sus GH13s and GH97s were inhibited similarly and the outer membrane α-amylase from Bo, BoGH13A_Sus_, was more inhibited by acarbose than SusG (**Table 2**). BoGH13A_Sus_ and SusG are required for growth on polysaccharides (32, 36, 51). The ability of Bo and Bt to grow on polysaccharides in the presence of acarbose concentrations higher than the BoGH13A_Sus_ and SusG IC_50_ values therefore suggests that these enzymes are not the primary acarbose target suggesting that the drug gets into the periplasm to disrupt maltooligosaccharide processing.

Because the deletion of BoGH97C_Sus_ and SusB eliminate the acarbose-induced growth lag to the same extent as *Δsus* deletions when cells are grown on maltose, we propose that these periplasmic enzymes drive the Bo and Bt susceptibility to acarbose, albeit to a lesser degree in Bo (**Fig. 6B**). However, we could not find a singular factor that explains the difference in growth phenotypes between Bo and Bt during acarbose exposure. We suspect that acarbose’s potent upregulation of *sus* might circumvent the lag when the primary target, the periplasmic GH97, is missing (**Fig. 8A**). Furthermore, we do not know which enzymes support BoΔBoGH97C_Sus_ and BtΔSusB growth on maltooligosaccharides (**Fig. 6A**). Although BoGH97D is likely upregulated by larger maltooligosaccharides (**Fig. 7**), the enzyme may not break down maltoheptaose optimally since BoΔBoGH97C_Sus_ lags compared to WT Bo when grown on this substrate (**Fig. 6A**). There are no obvious candidates from the RNAseq datasets comparing ΔSus and WT transcripts of Bo and Bt grown in maltose to explain how the strains lacking a Sus GH97 grow on maltooligosaccharides. Several SusC/D homologs are upregulated along with some GH18s and GH92s (**Tables S4** and **S5**). GH18 enzymes target chitin (β-linked N-acetylglucosamine) and GH92s are α-mannosidases (61). One or more of these enzymes may be promiscuous and target α-glucans, or α-glucans somehow get modified to become substrates for these enzymes. Despite the pivotal role that the Sus GH97s play in acarbose sensitivity, these enzymes do not explain the vastly different acarbose induced lag times between Bo and Bt (**Fig. 2B**).

An additional unexpected point of interaction between acarbose and the *Bacteroides* Sus is the outer membrane transporter. Acarbose likely competes with maltooligosaccharides for passage through the BoSusC/SusC transporter (**Fig. 4D**). BoΔBoGH13A_Sus_ and BtΔSusG exhibit an acarbose induced growth lag dependent on the size and shape of the oligo they are grown on (**Fig. 4**). We hypothesize that the lack of growth of cells on alpha-cyclodextrin (αCD) in the presence of acarbose is because acarbose is more readily transported through SusCD. However, this is somewhat surprising given that Bt SusD preferentially binds cyclic oligosaccharides over linear ones (34). SusD binding affinity may not be equivalent to SusC binding and transport. Additionally, tight binding by SusD may not be advantageous for growth if the “off” rate is too slow to pass the oligo through SusC. This seems unlikely given that both Bo and Bt grow well on αCD (**Fig. 4B**). Furthermore, acarbose still inhibits growth in BoΔBoSusCΔBoGH13A_Sus_ and BtΔSusCΔSusG strains, suggesting that the drug can enter the cell independent of BoSusC or SusC (**Fig. 4D**).

Finally, we identified a non-Sus GH97, BoGH97D, that is upregulated by maltose and acarbose in Bo (**Fig. 7**). Its Bt homolog, BtGH97H, is not upregulated in the same conditions (**Fig. 7D**). We initially hypothesized that BoGH97D could be responsible for Bo’s relative resistance to acarbose even if the enzyme itself is inhibited by acarbose (**Table 2**). For instance, BoGH97D could sequester acarbose away from BoGH97C_Sus_. However, Bt does not benefit when grown with acarbose while expressing BoGH97D constitutively (**Fig. 7E,F**). This may be because SusB is more abundant in the periplasm than BoGH97D so there is not enough of the latter to sequester acarbose away from SusB (**Fig. 8A**).

Though we have not determined a mechanism to explain Bo’s resistance to acarbose, two possibilities, among many, remain more likely. Bo upregulated its *sus* locus more strongly than Bt in maltose alone compared to glucose and in glucose supplemented with 50 µM acarbose compared to glucose alone (**Fig. 8A**). Given how potently, compared to maltose, acarbose upregulated *sus* in both organisms, it is possible that maltose is not the strongest *sus* inducer. Indeed, previous work on BtSus showed that maltoheptaose upregulates *sus* at lower concentrations than maltotriose (36). Taken together, BoSusR might sense maltooligosaccharides more efficiently than SusR and/or BoSusC mediated transport might be faster than SusC mediated transport. This would lead to faster buildup and increased concentrations of maltooligosaccharides in the Bo periplasm compared to the Bt periplasm.

Second, we initially checked that lysates of Bo and Bt grown on maltose do not break down acarbose (**Fig. S1B**). We wondered if the addition of acarbose to these growths would have any effect and therefore grew each organism in minimal media with maltose +/− 50 µM acarbose. Lysates from all conditions were active on maltose and maltoheptaose but not acarbose (**Fig. S12**). It is possible small amounts of acarbose were broken down to products not visible on the TLC, i.e., ones without a reducing end.

Lui et al predicted that acarbose targets intracellular enzymes in their work using fluorescently labelled maltodextrin to determine which bacteria are sensitive to acarbose (26). Species that were sensitive to acarbose induced growth inhibition still accumulated fluorescent maltodextrin signal in the periplasm in the presence of acarbose. Since this result was also observed in a particular strain lacking a BoGH13A_Sus_ or SusG homolog, the authors correctly predicted that acarbose inhibits periplasmic Sus enzymes, which we corroborate here coupled with previous data that SusB is inhibited by acarbose (42, 43). The inhibition of periplasmic enzymes and upregulation of *sus* by acarbose also explains why fluorescent maltodextrin accumulated in the periplasm (26). The authors correctly postulated that buildup of maltooligosaccharides in the periplasm due to enzyme inhibition would lead to more *sus* upregulation and therefore more maltodextrin transport. Our data support this observation, and we would add that acarbose also contributes to this effect because it upregulates *sus* (**Fig. 8A**) (26). In this case, more *sus* transcript would lead to more BoSusC/SusC transporter at the cell surface, thereby leading to increased maltooligosaccharide import and buildup in the periplasm due to intracellular enzyme inhibition by acarbose (26).

All told, acarbose interacts with Bo and Bt in predicted and unpredicted ways, as modelled in **Figs. 8B,C**. The results here are similar to what is observed in the *Escherichia coli* K12 mal system. Acarbose is transported by the maltoporin LamB but not appreciably metabolized by the MalQ or MalZ enzymes in *E. coli* (70). *E. coli* does not encode any GH97s in its genome while MalQ and MalZ are GH77 and GH13 family enzymes, respectively (61). Acarbose, however, is a much more potent inhibitor of *Bacteroides* growth than *E. coli* growth and only weakly upregulates the *mal* system in *E. coli*, unlike the robust *sus* upregulation we observed in Bo and Bt (70).

This newfound mechanistic understanding of how acarbose influences Bacteroidota growth, a phylum implicated in the etiology of diabetes, colorectal cancer, and colitis (71–75), will enable us to predict microbiome responsiveness to acarbose treatment. Furthermore, this work lays a foundation for the design of other xenobiotics with even stronger, or more tailored effects, than acarbose. Given the immense variability of the human gut microbiota, biochemical insight into species level interactions with acarbose is critical for repurposing this promising drug.

## Materials and Methods

### Bacterial Genetic Manipulation and Growth Conditions

*Bacteroides ovatus* (ATCC 8483, Bo) and *Bacteroides thetaiotaomicron* (VPI-5482, Bt) were routinely grown in tryptone-yeast extract-glucose (TYG) medium, minimal medium, or brain heart infusion (Becton Dickinson) agar supplemented with 10% horse blood (Colorado Serum Co.) (76, 77). For all growth comparisons and genome manipulations, BoΔ*tdk* and BtΔ*tdk* were used, unless otherwise noted, and are considered wild-type because they contain no *sus* mutations (34, 78). Numerous strains were previously generated via a previously described counter selectable allelic exchange method using a pExchange-*tdk* vector and are listed in **Supplementary Table 6** (34). To “delete” genes encoding the periplasmic Sus enzymes, a premature stop codon was introduced at Leu5 in both GH13s and Ser8 in both GH97s. pExchange plasmids were constructed with 750 bp flanks on either side of the premature stop codon. The gene for BoSusC (*bovatus_03807*) was deleted in both a WT and BoΔBoGH13A_Sus_ background. Full BoGH97C_Sus_ and SusB deletions were performed in a WT and BoΔBoGH13A_Sus_ and BtΔSusG background, respectively. These were subsequently used as parent strains to introduce the periplasmic GH97 from the opposite organism to generate BoΔBoGH97C_Sus_+SusB, BoΔBoGH97C_Sus_ΔBoGH13A_Sus_, BtΔSusB+BoGH97C_Sus_, and BtΔSusBΔSusG+BoGH97C_Sus_. The GH97 genes were incorporated in the native *sus* site using 750 bp flanks (*bovatus_03807* and *bovatus_03809*) to introduce SusB into the Bo genome and 750 bp flanks (*susC* and *SusA*) to introduce BoGH97C_Sus_ into the Bt genome.

The gene for BoGH97D (*bovatus_04772*) was cloned into a modified pNBU2 vector with a constitutively active promoter (sigma 70, *rpoD*) and complemented in a WT and BtΔSus background at the same tRNA^ser^ site to generate Bt::BoGH97D and BtΔSus::BoGH97D (69, 77).

All manipulated strains were screened for correct deletion or incorporation of the target gene by PCR-amplifying the expected inserts and sequencing them via Sanger Sequencing using the University of Michigan DNA sequencing core or Azenta Life Sciences. All strains are listed in **Supplementary Table 6**, and primers are listed in **Supplementary Table 7**.

For plate reader growths, strains were inoculated in TYG from freezer stocks and grown overnight at 37 °C in a Coy anaerobic chamber (85% N_2_/10% H_2_/5% CO_2_). Cells were back diluted 1:50 into MM + 5 mg/ml glucose, unless otherwise noted, and grown overnight. The following day, cells were pelleted and washed with 2x MM with no carbon source and diluted 1:100 into MM + 2.5 mg/ml carbohydrate +/− acarbose in parallel with a MM + 2.5 mg/ml glucose control. Substrates included maltose (Sigma), maltotriose (Carboexpert), maltoheptaose (Carbosynth), potato amylopectin (Sigma), bovine liver glycogen (Sigma), pullulan (Megazyme), soluble potato starch (Sigma), and α-cyclodextrin (Sigma). Cells were always mixed with a 2x solution of carbohydrate +/− acarbose to achieve the final 1:100 dilution. Kinetic growth experiments were done in 96-well plates in an anaerobic chamber at 37 °C on a BioTek Biostack plate-handler and Powerwave HT plate reader. Every 10 min, an OD_600_ of three replicates were recorded.

To calculate acarbose induced growth lag time, in hours, the average time to an OD_600_ of 0.3 in the -acarbose condition was calculated. This was subtracted from the time to OD_600_ of 0.3 in the +acarbose condition to calculate lag time.

### Gene Cloning and Site-Directed Mutagenesis for Heterologous Protein Expression

SusG and BoGH13A_Sus_ were previously cloned (38, 51). BoGH13B_Sus_ (locus tag *bovatus_03809*; NCBI: ALJ48414.1) and SusA (locus tag *bt_3704*; NCBI: AAO78809.1) were cloned using the Lucigen pET-ite system that introduced a TEV cleavage site before a 6x His tag. *Bovatus_03809* encodes a 684 amino acid protein whereas *bt_3704* encodes a 617 amino acid protein. The first annotation of the *B. ovatus* genome includes a BoGH13B_Sus_ gene (*bacova_03520*) that is 617 amino acids long. Therefore, we assumed this annotation is correct and that the *bovatus_03809* annotation is incorrect. We believe the true start site is at the currently annotated M68 because there is a 22 amino acid sequence following this start that is homologous to the SusA SPI signal peptide (62). SusA was cloned starting with T23 and BoGH13B_Sus_ was cloned starting with A90, hereafter referred to in terms of SusA numbering as A23.

BoGH97C_Sus_ (locus tag *bovatus_03808*; NCBI: ALJ48413.1), BoGH97D, (locus tag *bovatus_04772*; NCBI: ALJ49360.1), SusB (locus tag *bt_3703*; NCBI: AAO78808.1) and BtGH97H (locus tag *bt_4581*; NCBI: AAO79686.1) were cloned with the Lucigen pET-ite system without a TEV cleavage site since numerous attempts to TEV cleave these proteins were unsuccessful and previous work with SusB was done with enzyme retaining a 6x N-terminal His tag (42, 43). All GH97s here were cloned with an N-terminal 6x His tag.

BoGH13B_Sus_ and SusA genes were mutated to introduce a D to N mutation at position 331. This position aligns with the catalytic nucleophile in other GH13 enzymes. Site directed mutagenesis was performed via splicing by overlap-extension PCR (SOE-ing PCR).

All constructs were confirmed by Sanger Sequencing at the University of Michigan Sequencing Core or Azenta Life Sciences. Primers used during construct cloning are listed in **Supplementary Table 7**.

### Heterologous Protein Expression and Purification

Rosetta (DE3) pLysS cells were transformed with the appropriate construct and grown overnight in LB. 10 ml of LB per 500 ml of TB was used to inoculate TB medium the following morning (0.5 L for all GH97s and 1 L for all GH13s). Cells were grown to an OD_600_ of 0.6-0.7 at 37 °C and moved to room temperature to shake for 0.5 h before protein expression was induced with the addition of 0.5 mM isopropyl β-D-1-thiogalactopyranoside (IPTG). Cells were grown at room temperature (∼22 °C) for 18 h and harvested by centrifugation for 10 min at 10,000 x*g*. Cell pellets were stored at −80 °C. All proteins were purified according to (51) except that the GH97s were dialyzed against storage buffer (20 mM HEPES, 100 mM NaCl, pH 7.0) after the first purification because they were not TEV-cleaved.

### BoGH13B_Sus_ and SusA Size Exclusion Chromatography

0.5 ml of 10.6 mg/ml BoGH13B_Sus_ or 8.2 mg/ml SusA was applied to a HiPrep 16/60 Sephacryl S-200 HR (GE Healthcare) in 20 mM HEPES, 100 mM NaCl, pH 7.0 at a flow rate of 0.5 ml/min. BioRad gel filtration standards were reconstituted in 0.5 ml of the same buffer and run using the same protocol for comparison (bovine thyroglobulin [670 kDa], bovine γ-globulin [158 kDa], chicken ovalbumin [44 kDa], horse myoglobin [17 kDa], vitamin B12 [1.35 kDa]).

### α-glucan Breakdown by Cell Lysates

Bo, BoΔSus, Bt, BtΔSus, Bt::BoGH97D, and BtΔSus::BoGH97D were grown anaerobically in TYG from freezer stocks and back diluted into MM + 5 mg/ml maltose. Strains were back diluted to an OD_600_ of 0.2 in 10 ml of MM + 5 mg/ml maltose without acarbose and with 50 µM acarbose (for Bo and Bt only). Bacteria were grown to an OD_600_ of 0.7 and pelleted via centrifugation for 10 m at 5,000 rpm. Supernatant was decanted and cells were washed 2x in PBS. Cell pellets were flash frozen in liquid nitrogen and stored at −80 °C. Pellets were lysed in 1 ml of PBS via sonication. Half of the lysate was retained, and half was pelleted for 10 m at 13,000 x*g*. The pellet was discarded, and the clarified lysate was retained. Clarified and unclarified lysates were mixed 1:1 with 10 mg/ml maltose, maltoheptaose, or acarbose and incubated over night at 37 °C. Reaction products were analyzed via thin layer chromatography, described below.

### Thin Layer Chromatography

To assess oligosaccharide and polysaccharide break down by enzymes, thin layer chromatography (TLC) was performed as described in (51). 0.5 µM protein and 5 mg/ml of one of the following carbohydrates was used: maltose (Sigma), maltotriose (Carboexpert-BoGH13B_Sus_/SusA, Sigma-GH97s), maltotetraose (Carboexpert), maltopentaose (Carbosynth), maltohexaose (Carbosynth), maltoheptaose (Carboexpert), 6^3^-α-d-glucosyl-maltotriose (Megazyme), 6^3^-α-d-glucosyl-maltotriosyl-maltotriose (Megazyme), α-cyclodextrin (Sigma), β-cyclodextrin (Sigma), acarbose (Sigma), acarviosin (CarboSynth) D-panose (Sigma), isomaltose (Sigma), potato amylopectin (Sigma), bovine liver glycogen (Sigma), pullulan (Sigma), soluble potato starch (Sigma), and dextran (Sigma). All polysaccharides were autoclaved. A 20 mM HEPES, 100 mM NaCl (pH 7.0) buffer was used for BoGH13B_Sus_ and SusA while 100 mM maleate, 10 mM CaCl_2_ (pH 6.6) was used for BoGH97C_Sus_, BoGH97D, SusB, and BtGH97H according to conditions defined in (43). 2 µl of each reaction, as well as a no enzyme control, was spotted for separation and a 1 mg/ml solution of G1 – G7 in the appropriate reaction buffer was used as a ladder.

To resolve lysate reactions with maltose, maltoheptaose, or acarbose, 2 µl of each reaction, as well as a no lysate control, was spotted for separation and a 1 mg/ml solution of G1 – G7 in PBS was used as a ladder, along with acarbose and acarviosin in PBS as a comparison. TLC plate running and visualization conditions are described in (51).

### Enzyme Kinetics

BoGH13A_Sus_ and SusG inhibition was quantified using the EnzChek™ *Ultra* Amylase Assay Kit (Thermo). Briefly, the starch substrate in this assay is modified with a BODPY FL dye such that fluorescence is quenched. Upon digestion, the quenching is relieved and yields fluorescent fragments. The kit was used according to the manufacturer’s instructions except that a final concentration of 0.2 mg/ml fluorescent starch was used. 2x BoGH13A_Sus_ was preincubated +/− 2x 0.5 – 10 µM acarbose for 10 min and 2x SusG was preincubated +/− 2x 5 – 1000 µM acarbose for 10 min. Reactions were initiated via the mixing of 50 µl of 2x protein/buffer/acarbose and 50 µl of 2x starch for final concentrations of 25 nM enzyme, 0.2 mg/ml starch, and the previously indicated acarbose concentrations in 20 mM HEPES, 100 mM NaCl, pH 7.0. Reactions were performed in duplicate in black 96-well plates with clear bottoms (Corning) and monitored every 70 sec via excitation at 485 nm and emission at 528 nm in a Synergy H1 Plate Reader. Initial rates were calculated between 6 and 15 min. The 0 µM acarbose condition was treated as 100% activity and percent activity as a function of [acarbose] was graphed and analyzed in GraphPad Prism with a non-linear, [Inhibitor] vs. normalized response model to calculate an IC_50_.

BoGH13B_Sus_ and SusA Michaelis-Menten and inhibition kinetics were quantified using pNP-G6 (Sigma) in 20 mM HEPES, 100 mM NaCl, pH 7.0 Initial Michaelis-Menten parameters were determined with 1 nM enzyme in 25 – 750 µM pNP-G6 in duplicate. pNP release was monitored every minute at 405 nm in a Synergy H1 plate reader. Initial rates were calculated between 0 and 10 min. The *K*_M_ was subsequently used to inform inhibition conditions. Enzymes were pre-mixed with acarbose for 10 min and mixed 1:1 with pNP-G6 in duplicate. 60, 125, 250, 375, and 625 µM pNP-G6 was used with 0 – 480 nM acarbose and 2 nM enzyme. Inhibition was initially modelled in GraphPad Prism using a mixed model of inhibition. The subsequent low, but not zero, α value led us to do a final model using uncompetitive inhibition with an α*K*_i_ reported.

All GH97 (BoGH97C_Sus_, BoGH97D, SusB, and BtGH97H) kinetics were quantified using pNP-α-Glc (pNP-Glc, Sigma) in 100 mM maleate, 10 mM CaCl_2_, pH 6.6 according to (43). Initial Michaelis-Menten parameters for BoGH97C_Sus_ and SusB were used with 10 nM enzyme while 2 nM BoGH97D and BtGH97H was used with 10 – 500 µM pNP-Glc in duplicate. pNP release was monitored as described above and initial rates were determined between 0 and 10 min. The *K*_M_ was subsequently used to inform inhibition conditions. Enzymes were pre-mixed with acarbose for 10 min and mixed 1:1 with pNP-Glc in duplicate. 50, 100, 200, 300 and 500 µM pNP-Glc and 0 – 1000 µM acarbose was used. pNP release was monitored as described above and initial rates were determined between 0 and 10 min. GraphPad Prism was used to model competitive inhibition in all four enzymes to derive a *K*_i_.

### Structure of BoGH97C_Sus_ Bound to Acarbose

9 mg/ml of BoGH97C_Sus_ (using an estimated extinction coefficient of 156,268 M^−1^cm^−1^) with 10 mM acarbose was initially screened via sitting drop vapor diffusion at room temperature with an Art Robbins Gryphon LCP robot. Commercially available kits from Molecular Dimensions and Hampton were used for screening. Hanging drop refinement was carried out with a Molecular Dimensions Morpheus screen condition that included 100 mM NaHEPES/MOPS pH 7.5 (Thermo), 100 mM amino acids (from commercially available stock – includes L-Na-glutamate, racemic alanine, glycine, racemic lysine-HCl, and racemic serine), and 36% precipitant mix 4 (from commercially available stock 1:1:1 mix of racemic MPD, polyethylene glycol 1K, and polyethylene glycol 3350). Drops were a 1:1 mix of protein/acarbose:well solution. Suitable crystals were cryo-protected in a mix comprised of 70% well solution, 20% ethylene glycol, and 10% acarbose (final concentration of 10 mM) for 30 s before flash-freezing in liquid nitrogen. These crystals belonged to the orthorhombic space group *P*2_1_2_1_2_1_ with unit cell dimensions of *a* = 107.79 Å, *b* = 116.51 Å, *c* = 143.96 Å, α,β,γ = 90°. X-ray diffraction data were collected at the Advanced Photon Source Life Science Collaborative Access Team (LS-CAT) beamline 21 ID-G at Argonne National Laboratories. Data were processed and scaled in Xia2 with DIALS followed by molecular replacement in Phenix with PDB ID: 2ZQ0 (SusB) as a model (42, 79–81). Autobuild produced an initial structure with two monomers in the asymmetric unit (82). The model was manually adjusted in Coot followed by refinement in Refmac (83, 84). Acarbose geometry was validated using Privateer (85).

### Isothermal Titration Calorimetry

Acarbose binding to BoGH13B_Sus_ and SusA was assessed via isothermal titration calorimetry using a TA instruments standard volume ITC. All experiments were performed in triplicate at 25 °C using 25 µM BoGH13B_Sus_, BoGH13B_Sus_-D331, SusA, and SusA-D331N with 3.5 mM acarbose for the Bo proteins and 3 mM acarbose for the Bt proteins. A constant blank correction was used to account for the heat of dilution and all data were analyzed with an independent binding model using the manufacturer’s NanoAnalyze software.

### pNP-Glc Activity by Cell Lysates

Bo, BoΔSus, Bt, BtΔSus, Bt::BoGH97D, and BtΔSus::BoGH97D were grown in TYG overnight from freezers stocks and back diluted into MM+5 mg/ml maltose. The next morning, they were back diluted to an OD_600_ of 0.2 in 10 ml MM+5 mg/ml maltose and grown to an OD_600_ of 0.7. Cells were pelleted, washed 2x with PBS, pelleted, and flash frozen in liquid nitrogen and stored at −80 °C. Pellets were lysed in 1 ml of PBS via sonication. Debris was pelleted for 10 m at 13,000 x*g* and the clarified lysate was retained. Glucosidase activity was monitored with a 1:4 dilution of lysate in 1 mM pNP-Glc +/− 1 µM acarbose in PBS in duplicate. pNP release was monitored every minute at 405 nm in a Synergy H1 plate reader. Rates were calculated between 0 and 10 min.

### Activity Guided Fractionation

BoΔSus was first inoculated into TYG from a freezer stock and then back diluted, the next day, 1:100 in 50 ml of MM+5 mg/ml maltose. The following morning, all 50 ml were used to inoculate 1 L of MM+5 mg/ml maltose. Cells were grown to an OD_600_ of 0.75 and subsequently pelleted for 10 m at 22,000 *x*g at 4°C. The pellet was stored at - 80 °C for further processing. The BoΔSus pellet was resuspended in 80 ml of PBS and cells were lysed via sonication. The lysate was clarified via centrifugation at 30,000 *x*g for 30 m at 4 °C. 7.69g ammonium sulfate was added to 70 ml of clarified lysate to achieve 20% saturation and spun for 1 h at 4 °C. The mixture was centrifuged for 15 m at 10,000 *x*g at 4 °C. 20ml of the supernatant was saved for dialysis overnight against PBS. The process was repeated with the appropriate amount of solid ammonium sulfate to obtain 20 ml of mixtures at 30 and 50% and 10 ml at 70% saturation. All samples were dialyzed against 4 L of PBS at 4 °C overnight.

50 µl of each sample was mixed with 50 µl of 10 mM pNP-Glc to check for α-glucosidase activity. 50% saturation still afforded activity while 70% did not, so the protein sample from 50% saturation was used for downstream processing. The protein sample, in PBS, was applied to a 5 ml HiTrap Q FF column (Cytiva) preequilibrated in PBS. The column was washed and a 70 ml gradient from 100% 1x PBS (containing 137 mM NaCl and 2.7 mM KCl) to 100% PBS with extra salt in the same ratio (980 mM NaCl, 19.6 mM KCl). Only flow through fractions contained pNP-Glc activity so they were pooled, concentrated, and applied to a HiPrep 16/60 Sephacryl S-200 HR (GE) preequilibrated in PBS. Flow-through fractions were assessed for pNP-Glc activity and those with activity were pooled. They were applied to a 5 ml HiTrap SP FF column (Cytiva) in PBS and after dialyzing against 2-(N-morpholino)ethanesulfonic acid buffer at pH 6. No protein stuck to the column in either buffer. Active fractions were pooled and dialyzed against NaPO_4_ at either pH 5.1 or pH 8.4 to apply to a Q column equilibrated in the appropriate buffer. At pH 8.4, protein with pNP-Glc activity stuck to the column and was eluted in a gradient from 0 – 1 M NaCl in NaPO_4_. Active fractions were concentrated to 100 µl after dialysis overnight in 20 mM Bis-Tris propane (pH 8), 50 mM NaCl.

Samples were submitted to Proteomics Resource Facility at University of Michigan for analysis. Briefly, cysteines were reduced with 10 mM DTT (45° C for 30 min) and alkylated with 65 mM 2-Chloroacetamide, without light, for 30 min at room temperature. An overnight digestion with 1 µg sequencing grade modified trypsin was carried out at 37° C with constant mixing (ThermoMixer). The digestion was stopped by acidification and the peptides were desalted using SepPak C18 cartridges using the manufacturer’s protocol (Waters). Samples were then completely dried using a vacufuge. The resulting peptides were dissolved in 9 μl of 0.1% formic acid/2% acetonitrile solution. Two μls of the resulting peptide solution were resolved on a nano-capillary reverse phase column (EasySpray PepMap C18, 2 µm, 50 cm, #ES903, ThermoScientific) using a 0.1% formic acid/acetonitrile gradient at 300 nl/min over a period of 180 min. The eluent was directly introduced into a Q Exactive HF mass spectrometer (Thermo Scientific, San Jose CA) using an EasySpray source. MS1 scans were acquired at 60K resolution (AGC target=3×106; max IT=50 ms). Data-dependent collision induced dissociation MS/MS spectra were acquired on 20 most abundant ions following each MS1 scan (NCE ∼28%; AGC target 1×105; max IT 45 ms).

Proteins were identified by searching the data against the *Bacteroides ovatus* protein database (5499 entries) using Proteome Discoverer (v2.1, Thermo Scientific). Search parameters included MS1 mass tolerance of 10 ppm and fragment tolerance of 0.05 Da; two missed cleavages were allowed; carbamidomethylation of cysteine was considered fixed modification and oxidation of methionine, deamidation of asparagine and glutamine, variable modifications. False discovery rate (FDR) was determined using Percolator and proteins/peptides with an FDR of ≤1% were retained for further analysis. *GH97 Sequence Analysis*

Bo and Bt GH97 amino acid sequences according to the latest version of CAZy.org (61) were aligned in MegAlignPro using MAFFT (86, 87). A phylogenetic tree was made in MegAlignPro using the Neighbor-Joining algorithm and visualized in iTOL (88, 89). BoGH97C_Sus_, BoGH97D, SusB, and BtGH97H amino acid sequences were aligned and visualized using ClustalOmega on the EMBL-EBI server (90, 91).

### RNAseq Analysis

Bt, Bo, BtΔSus, and BoΔSus were grown anaerobically in TYG from freezer stocks and back diluted into MM+5 mg/ml maltose. Cells, in triplicate, were back diluted to an OD_600_ of 0.1 in 5 ml MM+5 mg/ml maltose and grown to an OD_600_ of 0.5. 10 ml of RNAprotect Bacteria Reagent (Qiagen) was added to cultures, incubated for 1 m, and cells were pelleted for 10 m at 3,500 rpm. Supernatant was removed and the pellets were stored at −80 °C until further processing.

Total RNA was purified according to (92) followed by treatment with DNase I (Invitrogen). RNA was repurified via standard sodium acetate/isopropanol precipitation and quantified using a NanoDrop (ThermoFisher). RNA, in water, was frozen at −80 °C. rRNA depletion, library preparation and sequencing were subsequently performed by SeqCenter (Philadelphia).

Raw reads were first trimmed and filtered via TrimGalore (version 0.6.6) and then aligned to the Bo or Bt protein-coding sequences via the DIAMOND aligner (version 2.0.6). Reads aligning to multiple protein coding references were adjudicated via FAMLI (https://github.com/Golob-Minot/FAMLI; (93)). The resultant specimen-gene-count matricies were outputted in anndata format. The entire workflow from raw reads to anndata count matricies was implemented in Nextflow, and available at https://github.com/jgolob/transcriptshot.

Differential gene expression was determined via Student T-tests, with Benjamini-Hochberg correction for false discovery. We used the following parameters to delineate significant up or down regulation: one mean ≥ 10 rpm; false discovery corrected students t-test p value ≥ 0.05; fold change ≥ 2.

### qPCR

Bo and Bt were grown in triplicate as described for the RNAseq analysis in MM + 5 mg/ml glucose or maltose +/− 50 µM acarbose. Total RNA was extracted from cells using an RNeasy kit (Qiagen), treated with DNase I (NEB), and reverse transcribed using Superscript III (Invitrogen) according to manufacturer’s protocols. The 16S rRNA gene-normalized transcript abundances were assayed on a Bio-Rad CFX Connect thermocycler using a custom qPCR master mix (94).

## Abbreviations

ACA: acarbose
Bo: *Bacteroides ovatus*
Bt: *Bacteroides thetaiotaomicron*
CBM: carbohydrate binding module
G2: maltose
G3: maltotriose
G4: maltotetraose
G5: maltopentaose
G6: maltohexaose
G7: maltoheptaose
GH: glycoside hydrolase
Glc: glucose
PUL: polysaccharide utilization loci
Sus: starch utilization system

---

**Supplementary Table 6.** Bacterial strains used in this study.

| Strain | Genotype | Reference |
| --- | --- | --- |
| Bo | BoΔtdk (BoΔbovatus_02336) | (78) |
| Bt | BtΔtdk (BtΔbt2275) | (34) |
| BoΔBoGH13A<sub>Sus</sub> | BoΔtdkΔbovatus_03803 | (51) |
| BtΔSusG | BtΔtdkΔbt3698 | (35) |
| BoΔBoSusC | BoΔtdkΔbovatus_3807 | This study |
| BtΔSusC | BtΔtdkΔbt3702 | (35) |
| BoΔBoSusCΔBoGH13A<sub>Sus</sub> | BoΔtdkΔbovatus_03803, Δbovatus_03807 | This study |
| BtΔSusCG | BtΔtdkΔbt3698Δbt3702 | (36) |
| BoΔBoGH13A<sub>Sus</sub>+SusG | BoΔtdkΔbovatus_03803+bt3698 | (51) |
| BtΔSusG+BoGH13A<sub>Sus</sub> | BtΔtdkΔbt3698+bovatus_03803 | (51) |
| BoΔBoGH13B<sub>Sus</sub> | BoΔtdk, bovatus_03809leu5 →stop | This study |
| BtΔSusA | BtΔtdk, bt3704leu5 →stop | This study |
| BoΔBoGH97C<sub>Sus</sub> | BoΔtdk, bovatus_03808ser8 →stop | This study |
| BtΔSusB | BtΔtdk, bt3703ser8 →stop | This study |
| BoΔBoGH97C<sub>Sus</sub>+SusB | BoΔtdkΔbovatus_03808+bt3703 | This study |
| BtΔSusB+BoGH97C<sub>Sus</sub> | BtΔtdkΔbt3703+bovatus_03808 | This study |
| BoΔSus | BoΔtdkΔbovatus_03803-bovatus03809 | N/A |
| BtΔSus | BtΔtdkΔbt3698-bt3704 | N/A |
| Bt::BoGH97D | BtΔtdk::bovatus_04772 | This study |
| BtΔSus::BoGH97D | BtΔtdkΔbt3698-bt3704::bovatus_04772 | This study |

**Supplementary Table 7.**
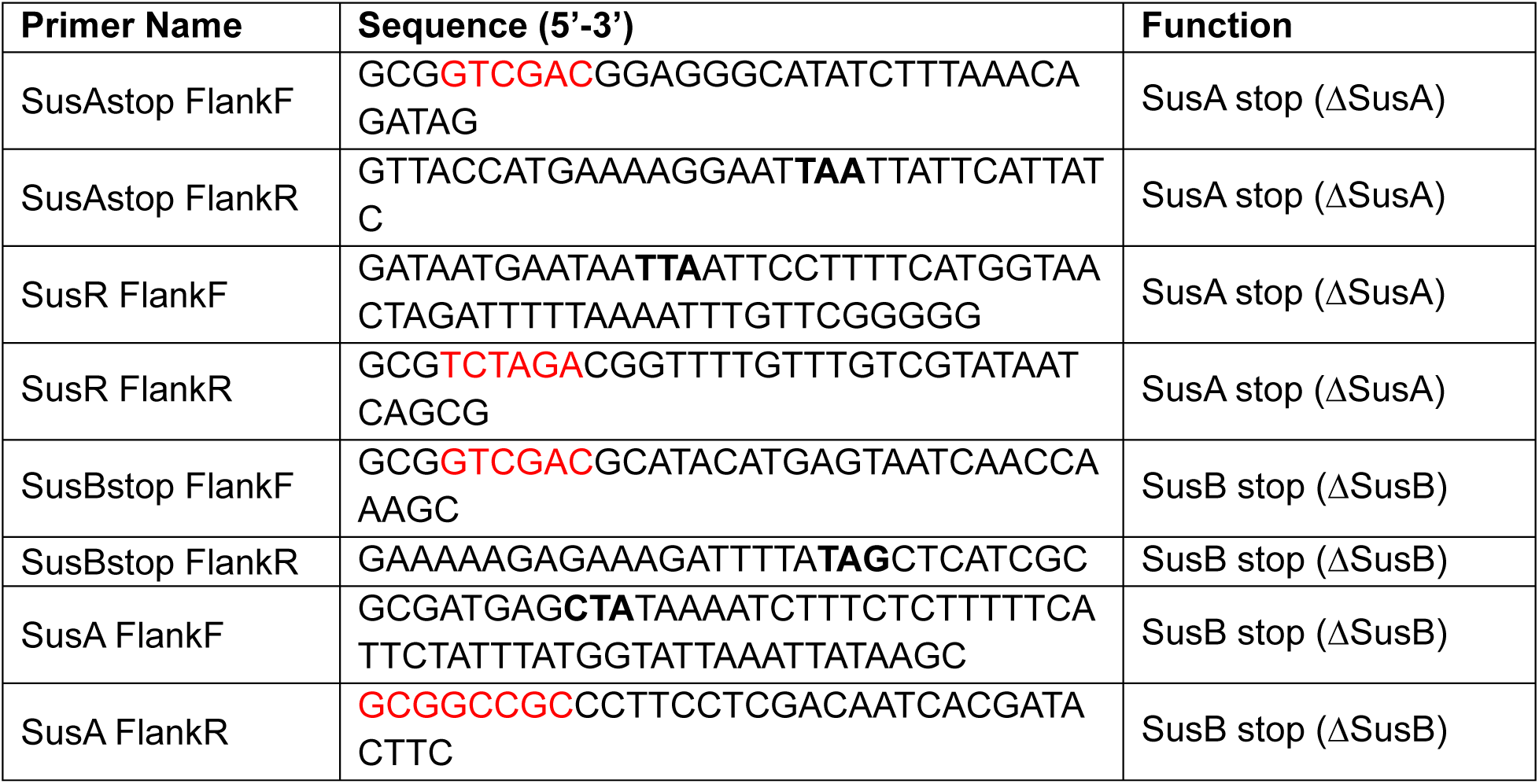
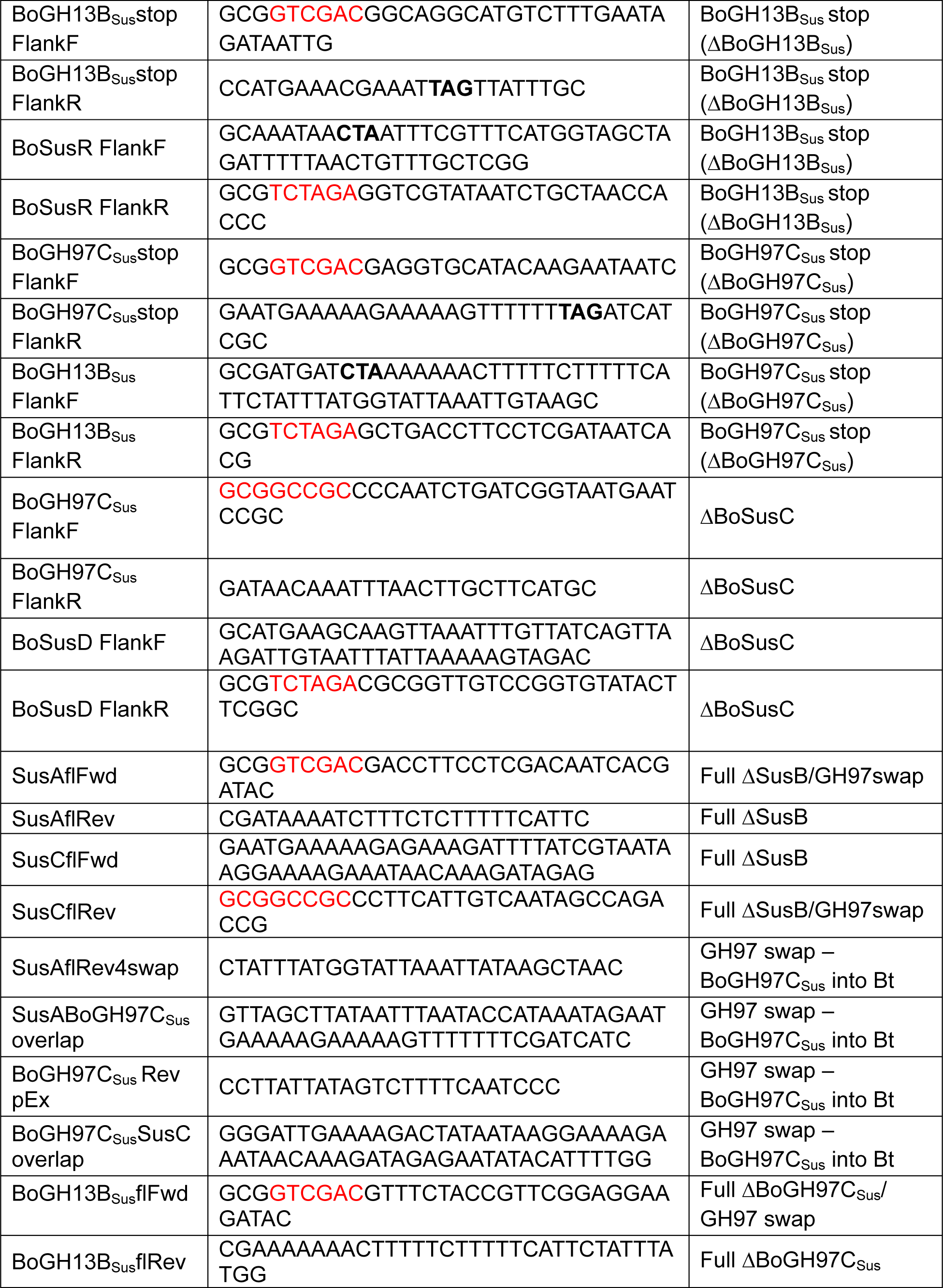
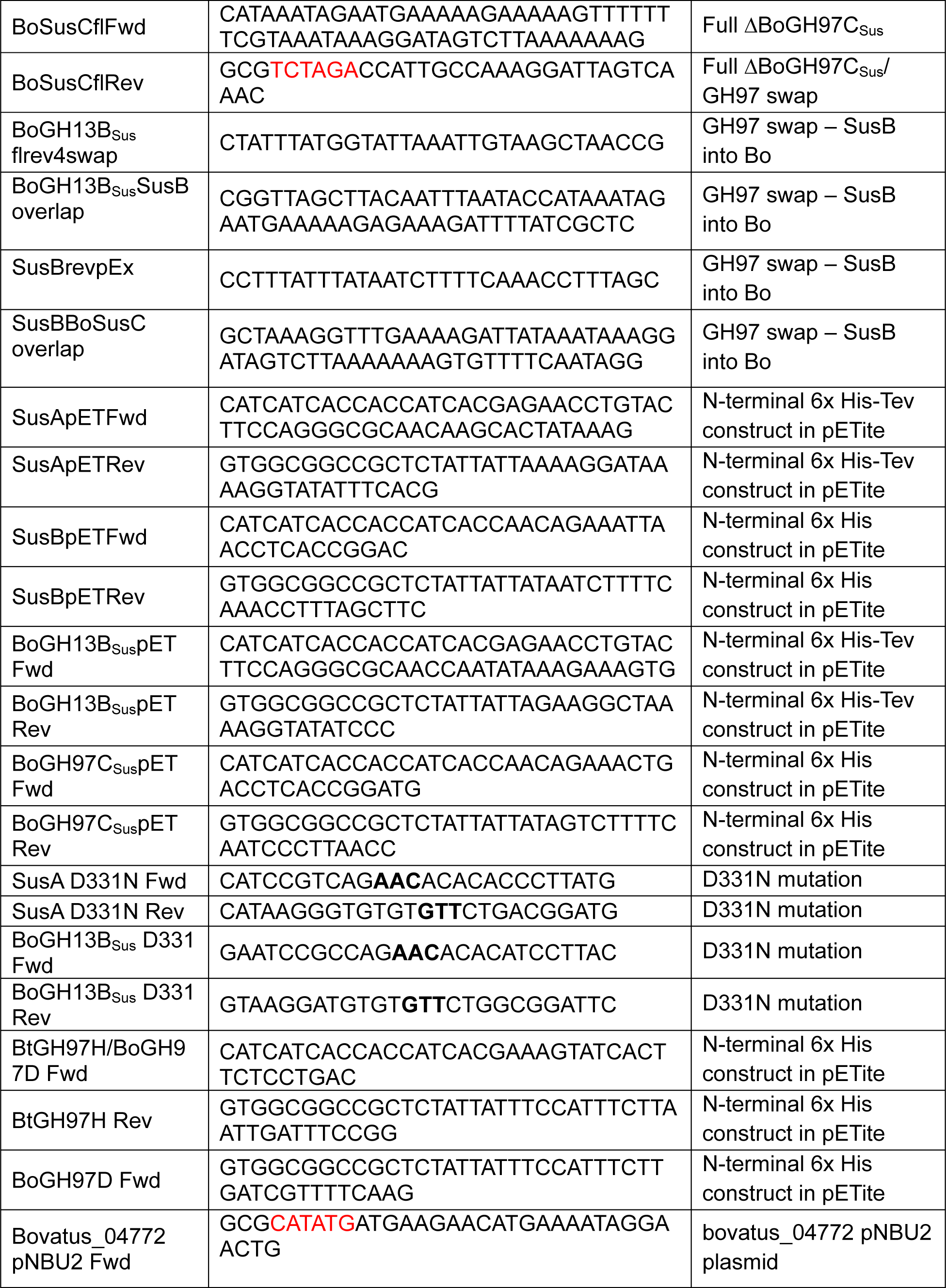
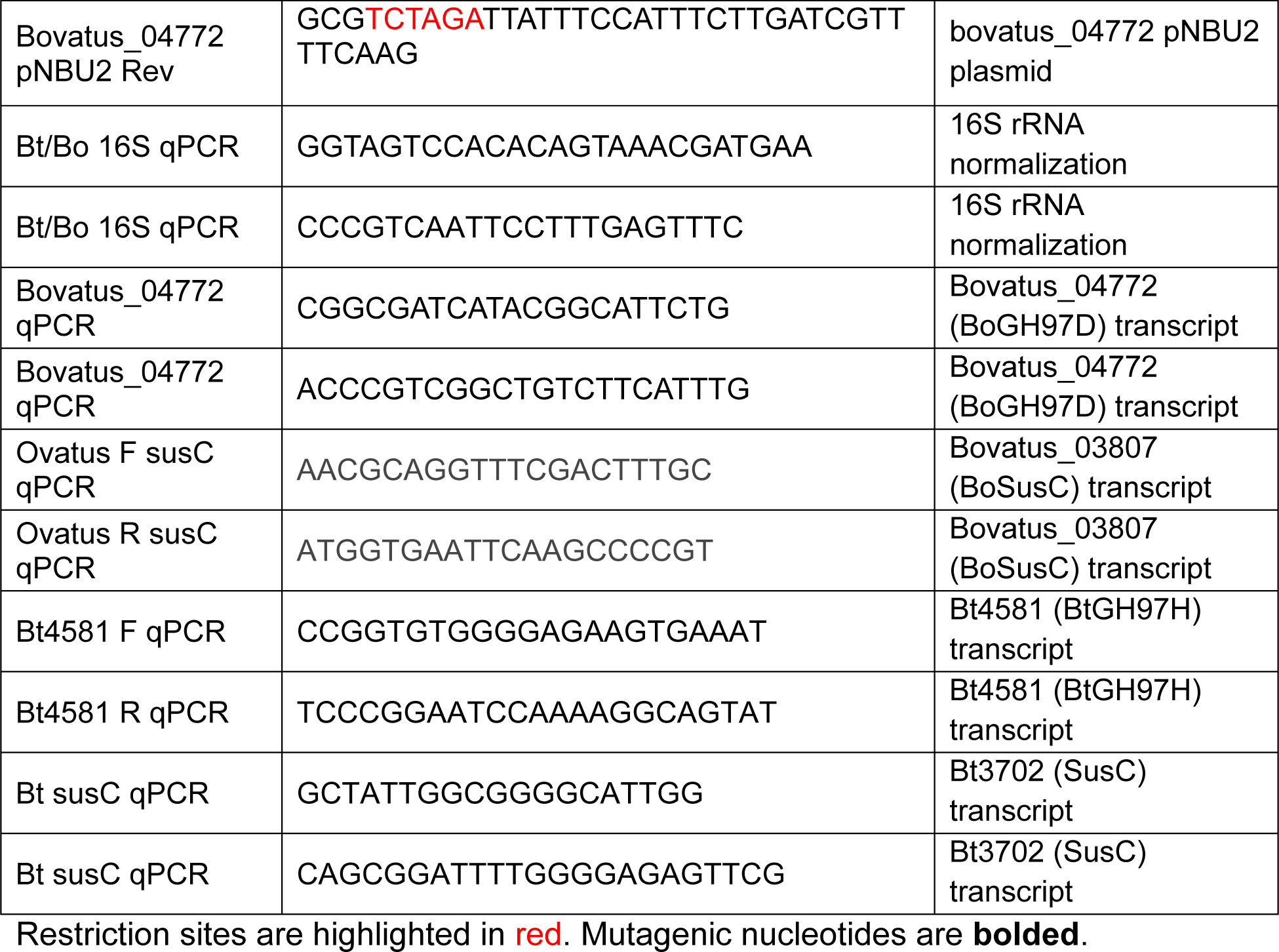
Oligonucleotide primers used in this study.

## Acknowledgements

We thank: Supriya Suresh Kumar who constructed the BtΔSus strain and Yunus E. Tuncil who constructed the BoΔSus strain; Darrell Cockburn for helpful suggestions on BoGH13B_Sus_ and SusA; Zdzislaw Wawrzak for help at the LS-CAT beamline; Venky Basrur for help at the University of Michigan Proteomics Resource Facility; members of the Koropatkin and Martens labs for helpful suggestions.

## Author Contributions

HAB: conceptualization, funding, investigation, project administration, supervision, data curation, data analysis, visualization, writing – original draft. ALM: investigation, data curation, data analysis. NAP: investigation, data curation, data analysis. AEH: investigation, data analysis. ECM: funding, resources. JLB: funding, investigation, data curation, data analysis. NMK: conceptualization, funding, data curation, project administration, supervision, editing.

## Data and Material Availability

RNAseq data was deposited under the NIH sequence read archive BioProject PRJNA1107240. Proteomics data was deposited in the ProteomeXchange Consortium via the Proteomics IDEntifications Database (PRIDE) partner repository with the dataset identifier of PXD052070 (95). The BoGH97C_Sus_ – acarbose structure was deposited with the RCSB Protein Databank with a PDB ID of: 9BS5. All plasmids, proteins, bacterial strains, and other reagents generated for this work will be made freely available to researchers using them for non-commercial reasons.

