## Supplemental information for "Acarbose Impairs Gut *Bacteroides* Growth by Targeting Intracellular GH97 Enzymes"

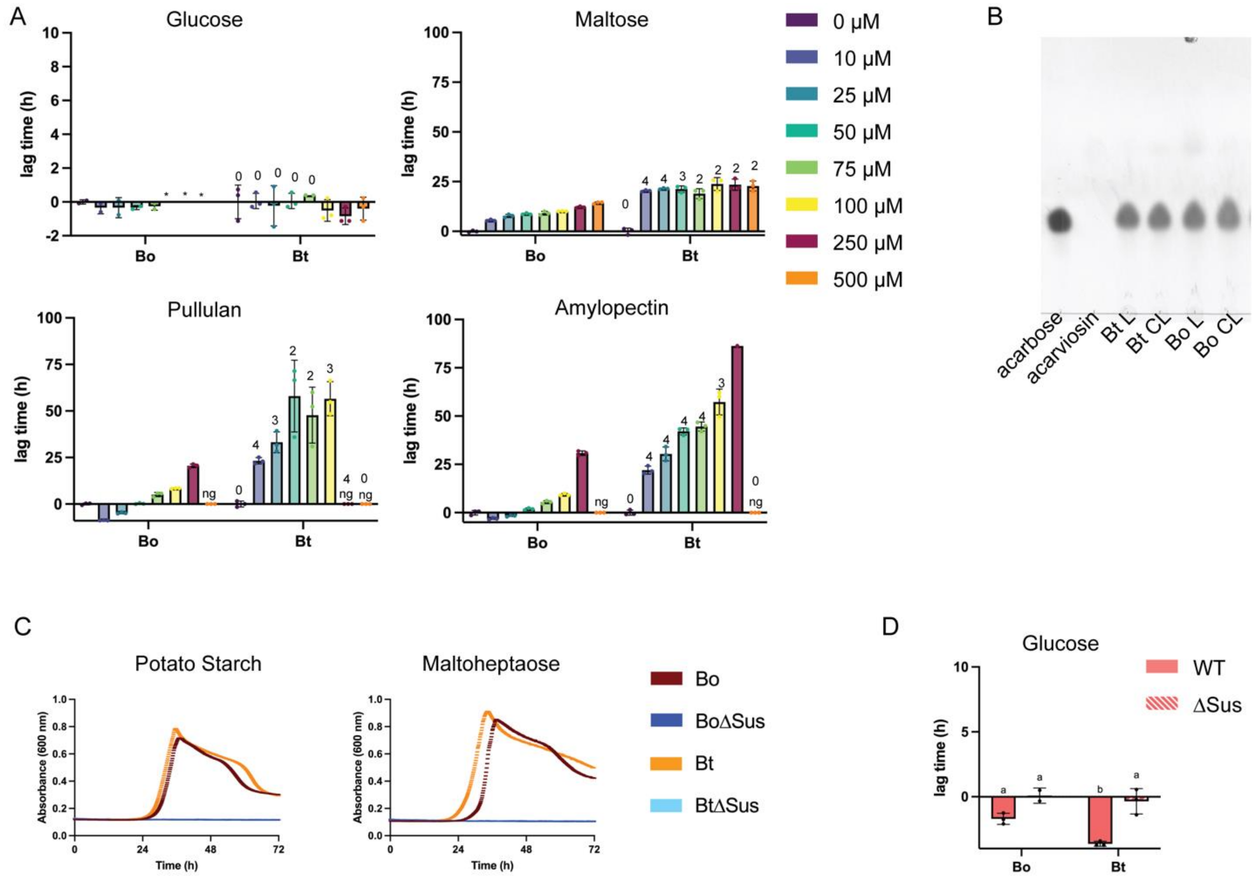


**Supplementary Figure 1. Bt is more susceptible to acarbose induced growth inhibition than Bo.** A) Bo and Bt were pre-grown in minimal media (MM) with glucose and back diluted into MM + 2.5 mg/ml of the indicated carbon sources with and without 10 – 500 μM acarbose. The difference in time to $OD_{600}$ of 0.3 between the treated and untreated conditions are graphed. Bo and Bt with the same treatment were compared using an unpaired, two-tailed Student's *t* test. 1: $p \leq 0.05$; 2: $p \leq 0.01$; 3: $p \leq 0.001$. 4: $p \leq 0.0001$. ng = no growth. Technical difficulties with the plate reader did not allow us to calculate an acarbose induced lag time at an $OD_{600}$ of 0.3 for conditions marked with *. B) Bo and Bt were grown in MM + maltose to an $OD_{600}$ of 0.7. Cells were pelleted and washed with PBS and sonicated to release cellular contents. L = lysate. CL = clarified lysate. Lysate and clarified lysate were incubated overnight at 37 °C with 5 mg/ml acarbose. 5 mg/ml acarbose and acarviosin in PBS alone were run out on a thin layer chromatography (TLC) plate as reference points. The acarviosin is O-methylated at the reducing end and is thus not visible because it cannot react with the visualizing agent, orcinol, which reacts with reducing ends. The faint bands in the experimental lanes are likely left over maltose from the MM. C) WT and ΔSus strains of Bo and Bt were pre-grown as described in A and back diluted into MM + 2.5 mg/ml amylopectin or maltoheptaose. D) WT and ΔSus strains of Bo and Bt were pre-grown as described in A and back diluted into MM + 2.5 mg/ml glucose +/- 50 μM acarbose. The difference in time to $OD_{600}$ of 0.3 between the treated and untreated conditions are graphed. Statistical analyses were performed using a two-way ANOVA with a cutoff of $p \leq 0.05$. Condition(s) with the same letter were not statistically different from one another.

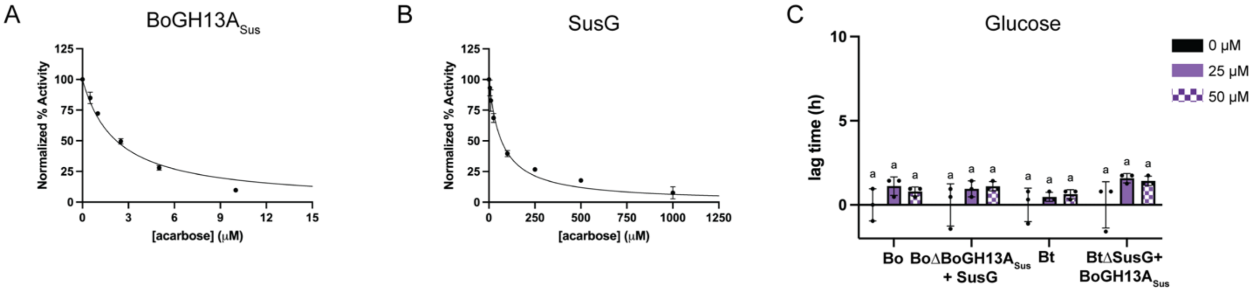


**Supplementary Figure 2. Outer membrane amylases BoGH13A$_{Sus}$ and SusG are not the source of the acarbose lag phenotype.** A,B) BoGH13A$_{Sus}$ and SusG starch breakdown in the presence of various acarbose concentrations was assessed using an EnzChek™ *Ultra* Amylase Assay Kit from Thermo. 25 nM enzyme and 0.2 mg/ml fluorescent starch was used. Activity in the absence of acarbose was set to 100% and percent activity of this was graphed vs. acarbose concentration to calculate $IC_{50}$ values, reported in Table 1. C) Because BoGH13A$_{Sus}$ does not optimally complement BtΔSusG growth on starch, these growths were performed with bacteria pre-grown on minimal media (MM) + 5 mg/ml maltose to induce *sus* expression, then inoculated into MM + 2.5 mg/ml glucose in the indicated acarbose concentrations as controls. Acarbose induced lag times are shown. Statistical analyses were performed using a two-way ANOVA using a cutoff of $p \leq 0.05$. Conditions with the same letter(s) were not significantly different from one another.

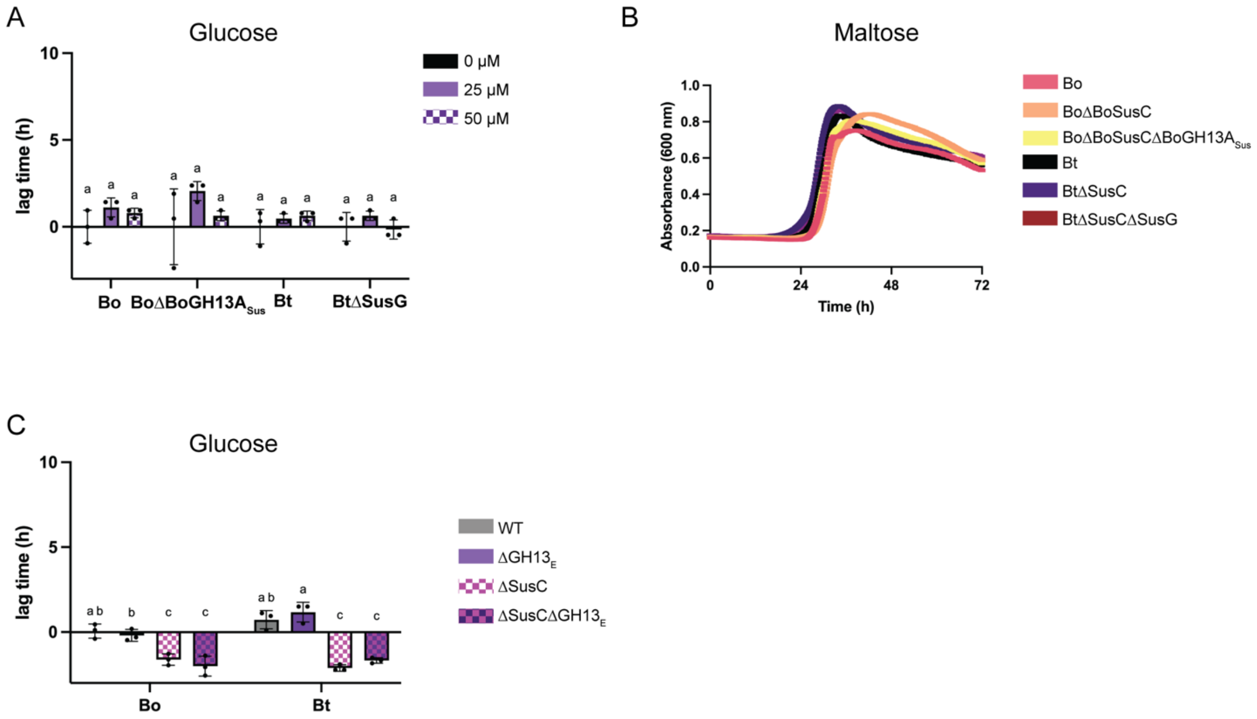


**Supplementary Figure 3. Acarbose likely competes with maltooligosaccharides for transport through BoSusC and SusC.** A) Bacteria were pre-grown in minimal media (MM) with glucose and back diluted in MM + 2.5 mg/ml glucose with and without the indicated acarbose concentrations. Acarbose induced lag times are shown. B) Bacteria were pre-grown in MM with glucose and back diluted in MM + maltose. C) Bacteria were pre-grown in minimal media (MM) with glucose and back diluted in MM + 2.5 mg/ml glucose with and without 50 μM acarbose. $\Delta$GH13$_E$ (GH13 extracellular) corresponds to Bo$\Delta$BoGH13A$_{Sus}$ and Bt$\Delta$SusG. Acarbose induced lag times are shown. Statistical analyses in A and C were performed with a two-way ANOVA. Conditions with the same letter(s) were not significantly different from one another. A cutoff of $p \leq 0.05$ was used.

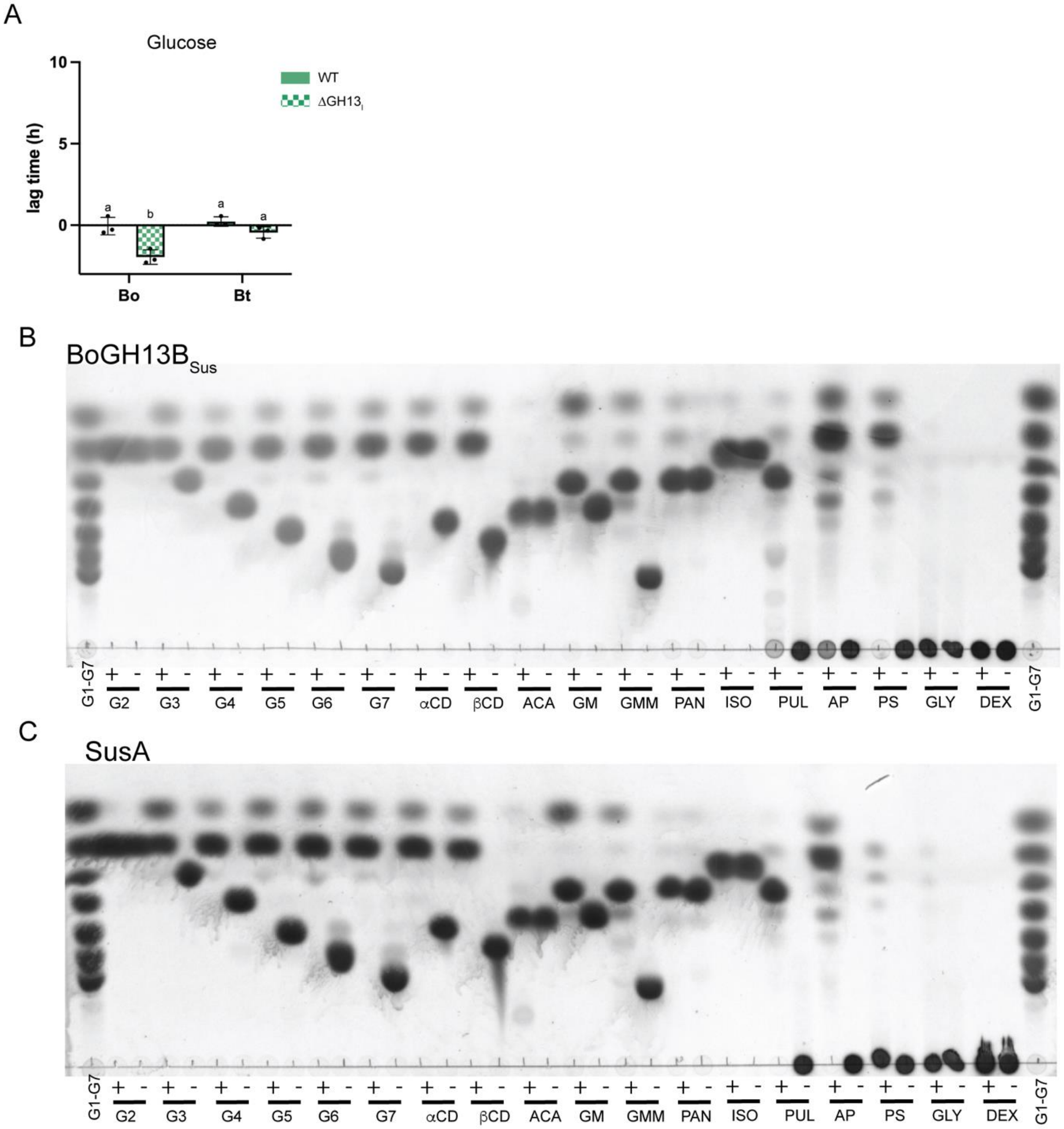


**Supplementary Figure 4. BoGH13B$_{Sus}$ and SusA do not underpin the different Bo and Bt acarbose phenotypes and have nearly identical substrate preferences.** A) Bo and Bt were pre-grown in minimal media (MM) with glucose and back diluted into MM + 2.5 mg/ml glucose with or without 50 µM acarbose. $\Delta$GH13$_I$ (GH13 intracellular)

corresponds to Bo$\Delta$BoGH13B$_{Sus}$ and Bt$\Delta$SusA. Acarbose induced growth lags are displayed and statistical analyses were performed with a two-way ANOVA. Conditions with the same letter(s) were not significantly different from one another. A cutoff of $p \leq 0.05$ was used. B-C) 500 nM of the indicated enzyme was incubated overnight with 5 mg/ml of the following carbohydrates: G2 – maltose; G3 – maltotriose; G4 – maltotetraose; G5 – maltopentaose; G6 – maltohexaose; G7 – maltoheptaose; $\alpha$CD – alpha-cyclodextrin; $\beta$CD – beta-cyclodextrin; ACA – acarbose; GM – $6^3$-$\alpha$-D-glucosyl-maltotriose; GMM – $6^3$-$\alpha$-D-glucosyl-maltotriosyl-maltotriose; PAN – D-panose; ISO – isomaltose; PUL – pullulan; AP – potato amylopectin; PS – potato starch; GLY – glycogen; DEX – dextran. All polysaccharides were autoclaved to get them into solution. + = with enzyme. - = no enzyme control.

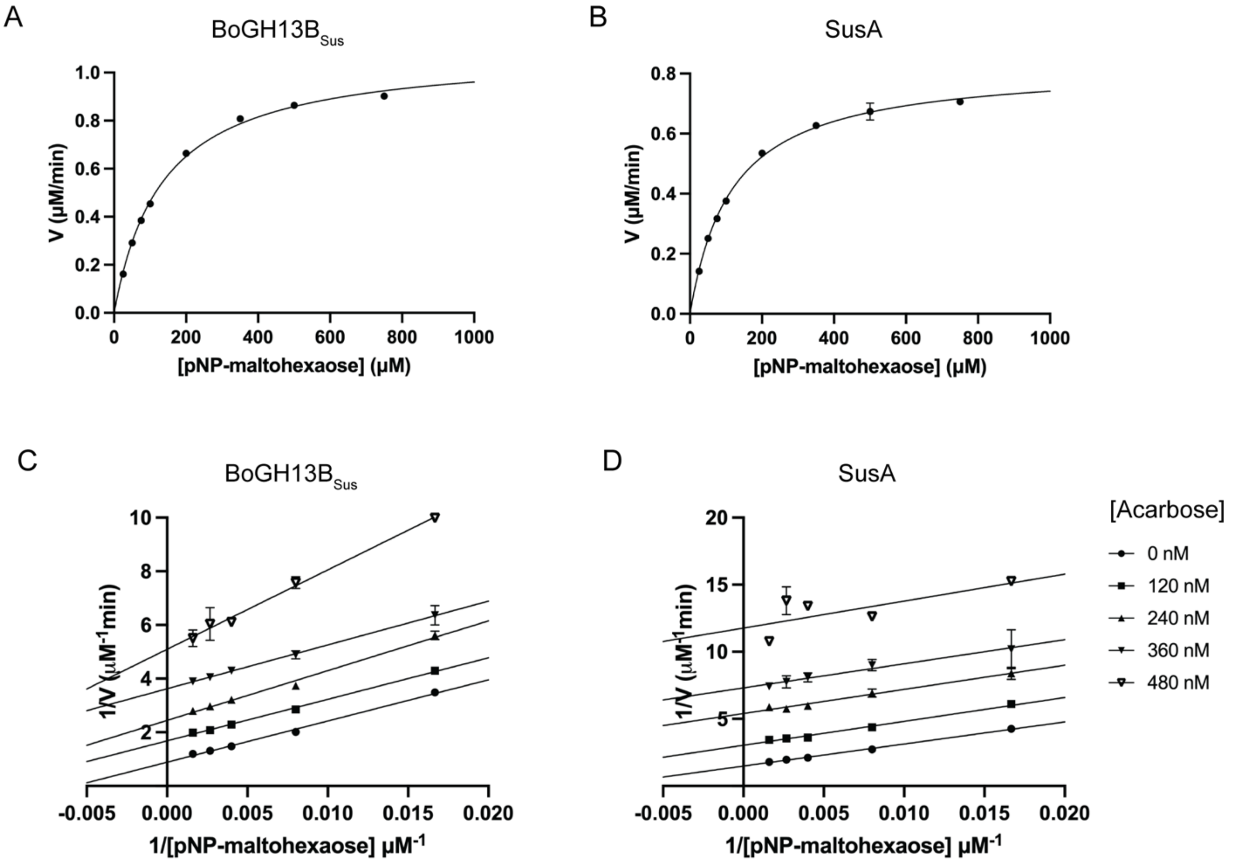


**Supplementary Figure 5. BoGH13B$_{Sus}$ and SusA have similar catalytic efficiencies and are inhibited similarly by acarbose.** A,B) Enzymes were incubated for 10 min in various concentrations of pNP-maltohexaose in duplicate. Initial rate as a function of substrate concentration is shown. C,D) Enzymes were pre-incubated for 10 min with acarbose before mixing with various pNP-maltohexaose concentrations. Initial rates were recorded and used to make the double reciprocal Lineweaver-Burk plots shown. Michaelis-Menten and inhibition parameters can be found in Table 1. 1 nM of each enzyme was used for both experiments.

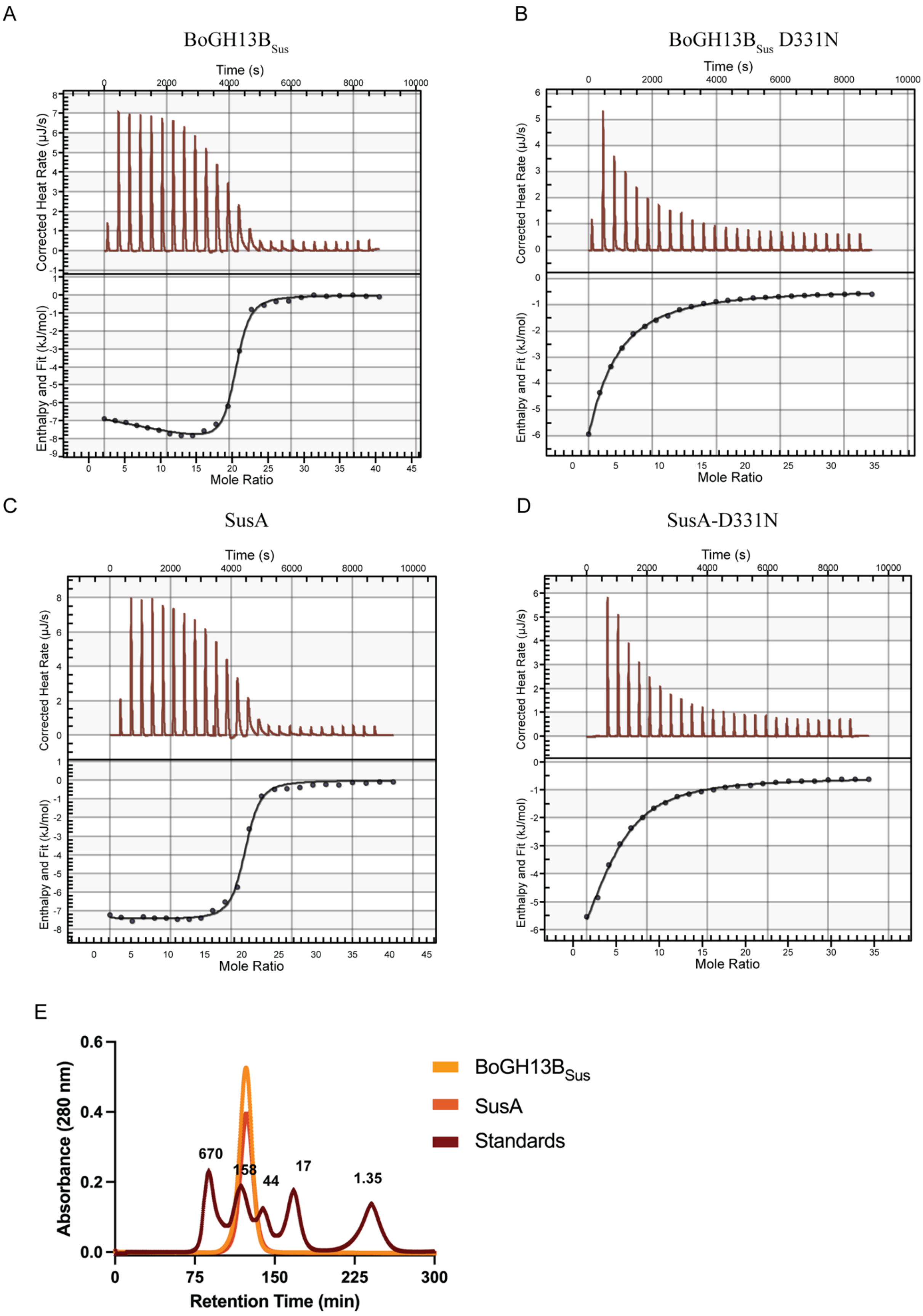
A
BoGH13BSus
Time (s)
Corrected Heat Rate (µJ/s)
Enthalpy and Fit (kJ/mol)
Mole Ratio
B
BoGH13BSus D331N
Time (s)
Corrected Heat Rate (µJ/s)
Enthalpy and Fit (kJ/mol)
Mole Ratio
C
SusA
Time (s)
Corrected Heat Rate (µJ/s)
Enthalpy and Fit (kJ/mol)
Mole Ratio
D
SusA-D331N
Time (s)
Corrected Heat Rate (µJ/s)
Enthalpy and Fit (kJ/mol)
Mole Ratio
E
Absorbance (280 nm)
Retention Time (min)
670
158
44
17
1.35
BoGH13BSus
SusA
Standards

**Supplementary Figure 6. BoGH13B$_{Sus}$ and SusA are both dimers that bind tightly to acarbose in the absence of substrate.** A-D) 25 μM of each enzyme was assessed for binding to 3.5 mM (Bo enzymes) or 3 mM (Bt enzymes) using a standard volume isothermal titration calorimeter from TA Instruments. Experiments were performed in triplicate with the average $K_D$ and n values being reported in Supplementary Table 1. Representative curves are displayed for each condition. E) BoGH13B$_{Sus}$ and SusA were applied to a HiPrep 16/60 Sephacryl S-200 HR column. Monomers of each enzyme are expected at ~69 kDa and dimers at ~138 kDa. Comparing the retention times of each enzyme to a standard curve of the logarithm of the retention time of the standards led to an estimated molecular weight of 125 kDa for BoGH13B$_{Sus}$ and 121 kDa for SusA, consistent with dimer formation.

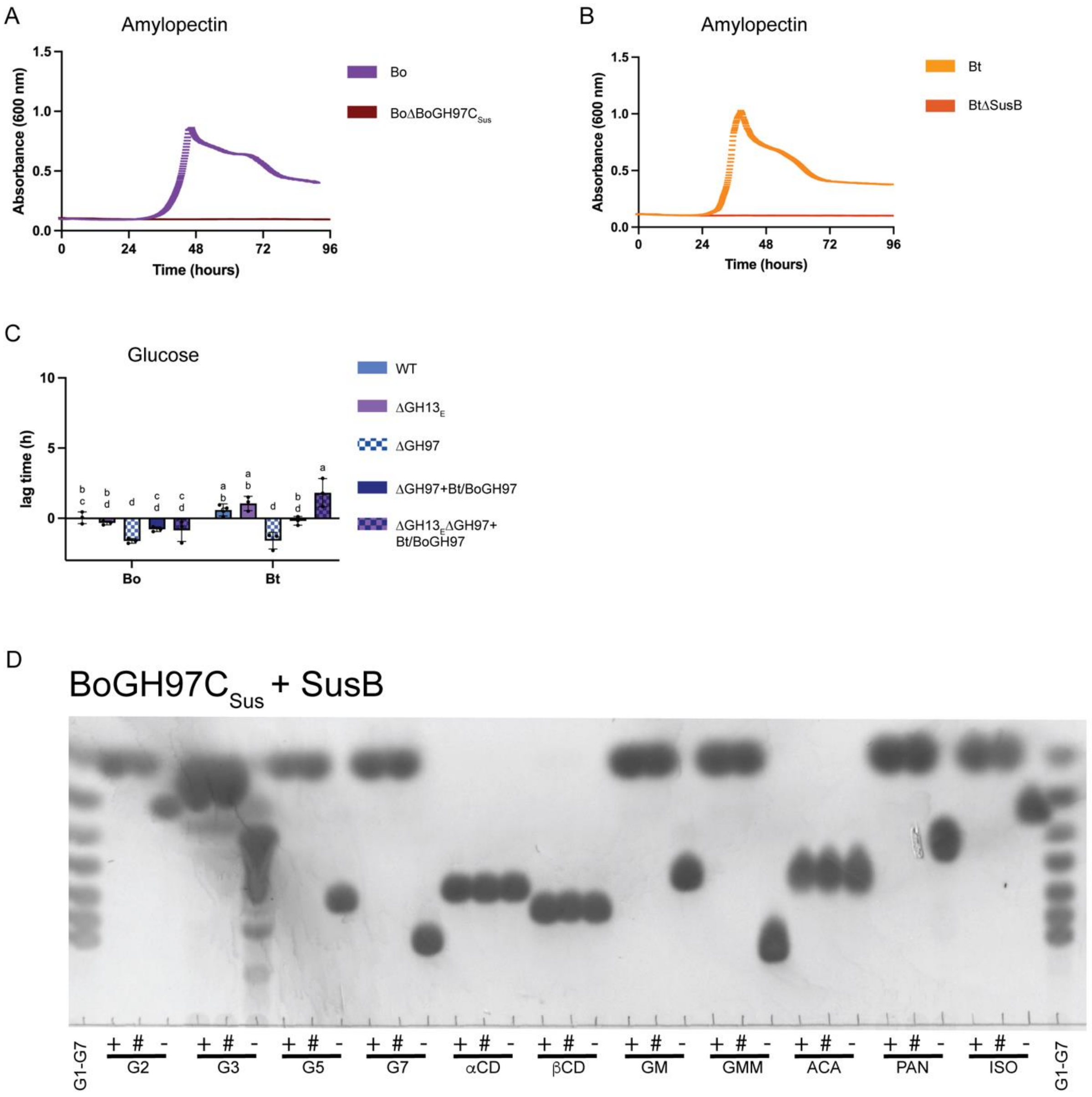


**Figure 7. Periplasmic Sus GH97 enzymes are the primary acarbose target but do not explain the different Bo and Bt phenotypes in acarbose.** A,B) Bacteria were pre-grown on minimal media (MM) with glucose and back diluted into MM + 2.5 mg/ml potato amylopectin. C) Bacteria were pre-grown in MM with glucose and back diluted into MM + 2.5 mg/ml glucose with our without 50 µM acarbose. ΔGH13$_E$ (GH13 extracellular) corresponds to BoΔBoGH13A$_{Sus}$ and BtΔSusG. ΔGH97 corresponds to BoΔBoGH97C$_{Sus}$ and BtΔSusB. Acarbose induced growth lags are displayed and

statistical analyses were performed with a two-way ANOVA. Conditions with the same letter(s) were not significantly different from one another. A cutoff of $p \leq 0.05$ was used. D) 500 nM of the indicated enzyme was incubated overnight with 5 mg/ml of the following carbohydrates: G2 – maltose; G3 – maltotriose; G5 – maltopentaose; G7 – maltoheptaose; αCD – alpha-cyclodextrin; βCD – beta-cyclodextrin; GM – $6^3$-α-D-glucosyl-maltotriose; GMM – $6^3$-α-D-glucosyl-maltotriosyl-maltotriose; ACA – acarbose; PAN – D-panose; ISO – isomaltose. + = BoGH97C$_{Sus}$. # = SusB. - = no enzyme control.

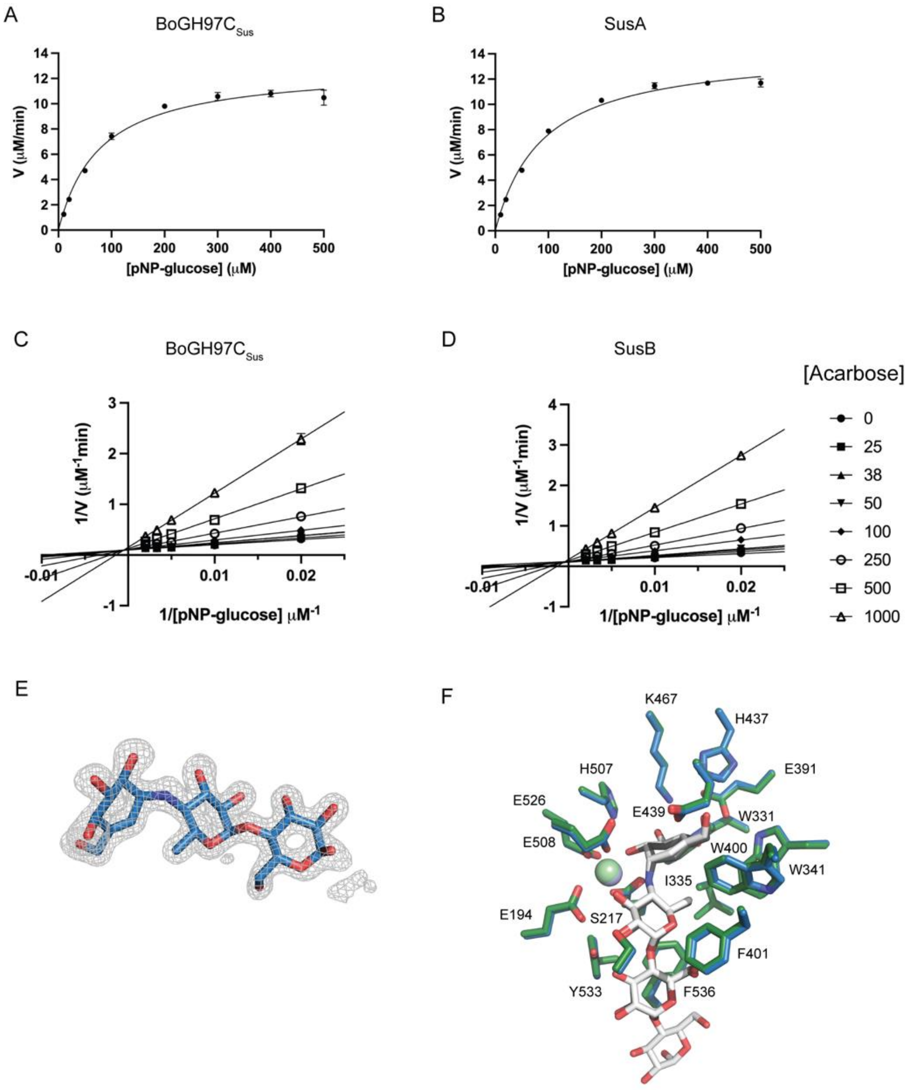


**Supplementary Figure 8. BoGH97C$_{Sus}$ and SusB are similarly inhibited by acarbose.** A,B) Enzymes were incubated for 10 min in various concentrations of pNP-glucose in duplicate. Initial rate as a function of substrate concentration is shown. C,D) Enzymes were pre-incubated for 10 min with acarbose before mixing with various pNP-glucose concentrations. Initial rates were recorded and used to make the double reciprocal Lineweaver-Burk plots shown. Michaelis-Menten and inhibition parameters can be found in Table 1. 10 nM of each enzyme was used for both experiments. E) $F_o$-$F_c$ density for acarviosin-glucose bound to chain A in the BoGH97C$_{Sus}$ model. While there was some density for a second glucose, an entire acarbose molecule could not be modelled in accurately. Density was contoured to 3$\sigma$. F) Comparison of acarbose bound to SusB and acarviosin-glucose bound to BoGH97$C_{Sus}$. The acarviosin-glucose sticks are black and BoGH97$C_{Sus}$ side chains in blue. The acarbose sticks are white and SusB side chains are in green (PDB ID: 2ZQ0, (1). Bound $Ca^{2+}$ is blue (BoGH97$C_{Sus}$) and green (SusB). The superpose command in PyMOL was used to overlay the structures with a root mean squared deviation (rmsd) of 0.46 Å over all atoms. E and F were rendered in PyMOL (2).

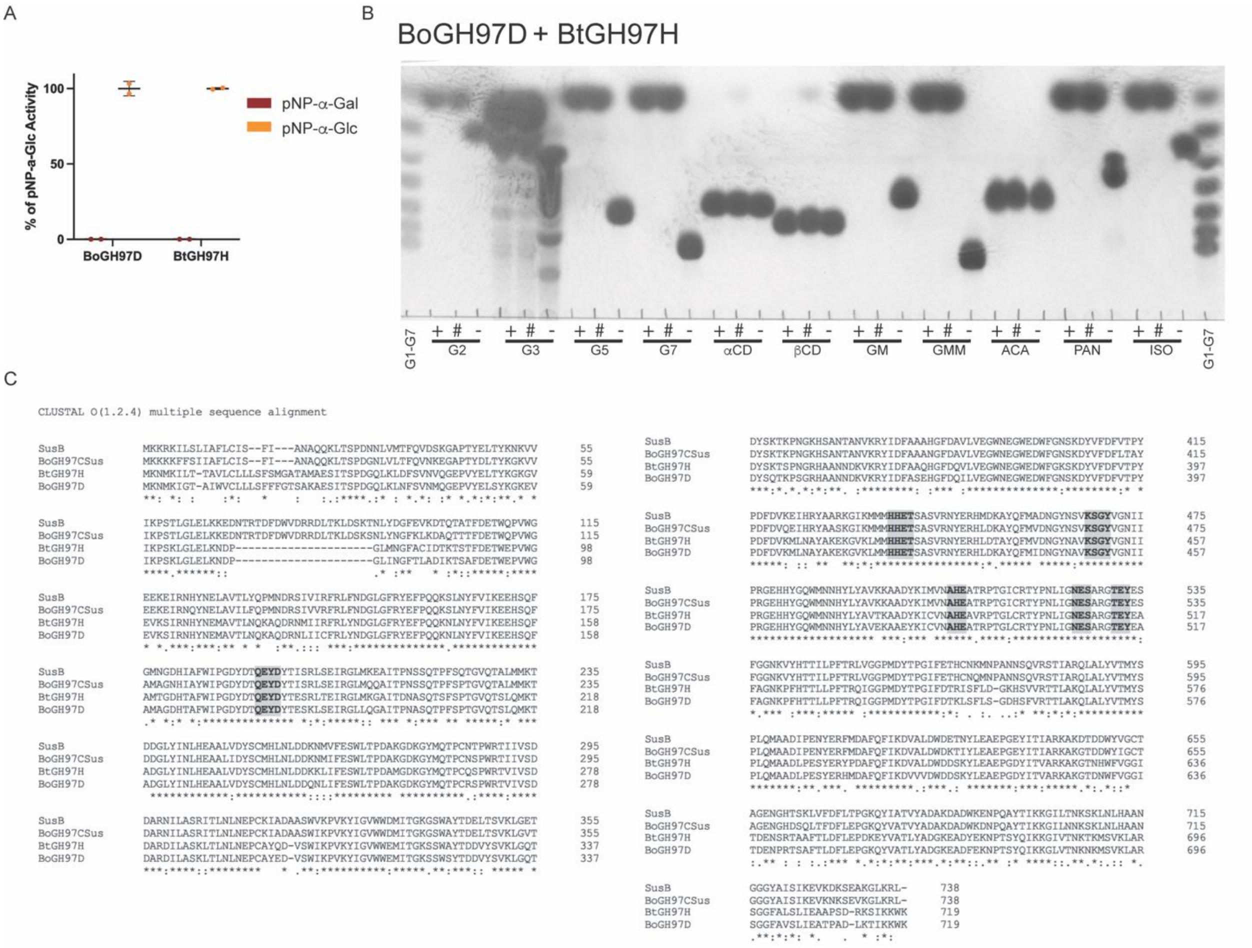


**Supplementary Figure 9. BoGH97D and BtGH97H are likely inverting α-glucosidases/glucoamylases.** A) 10 nM of each enzyme was incubated with 1 µM pNP-α-galactose (pNP-α-Gal) or pNP-α-glucose (pNP-α-Glc) and initial rates were recorded over ten minutes. Percent of pNP-α-Glc activity is graphed. B) 500 nM of the indicated enzyme was incubated overnight with 5 mg/ml of the following carbohydrates: G2 – maltose; G3 – maltotriose; G5 – maltopentaose; G7 – maltoheptaose; αCD – alpha-cyclodextrin; βCD – beta-cyclodextrin; GM – $6^3$-α-D-glucosyl-maltotriose; GMM – $6^3$-α-D-glucosyl-maltotriosyl-maltotriose; ACA – acarbose; PAN – D-panose; ISO – isomaltose. + = BoGH97C$_{Sus}$. # = SusB. - = no enzyme control. C) The indicate enzyme amino acid sequences were aligned using ClustalOmega in the EMBL-EBI server (3).

Amino acid signatures of inverting GH97s according to (4) are highlighted in grey.

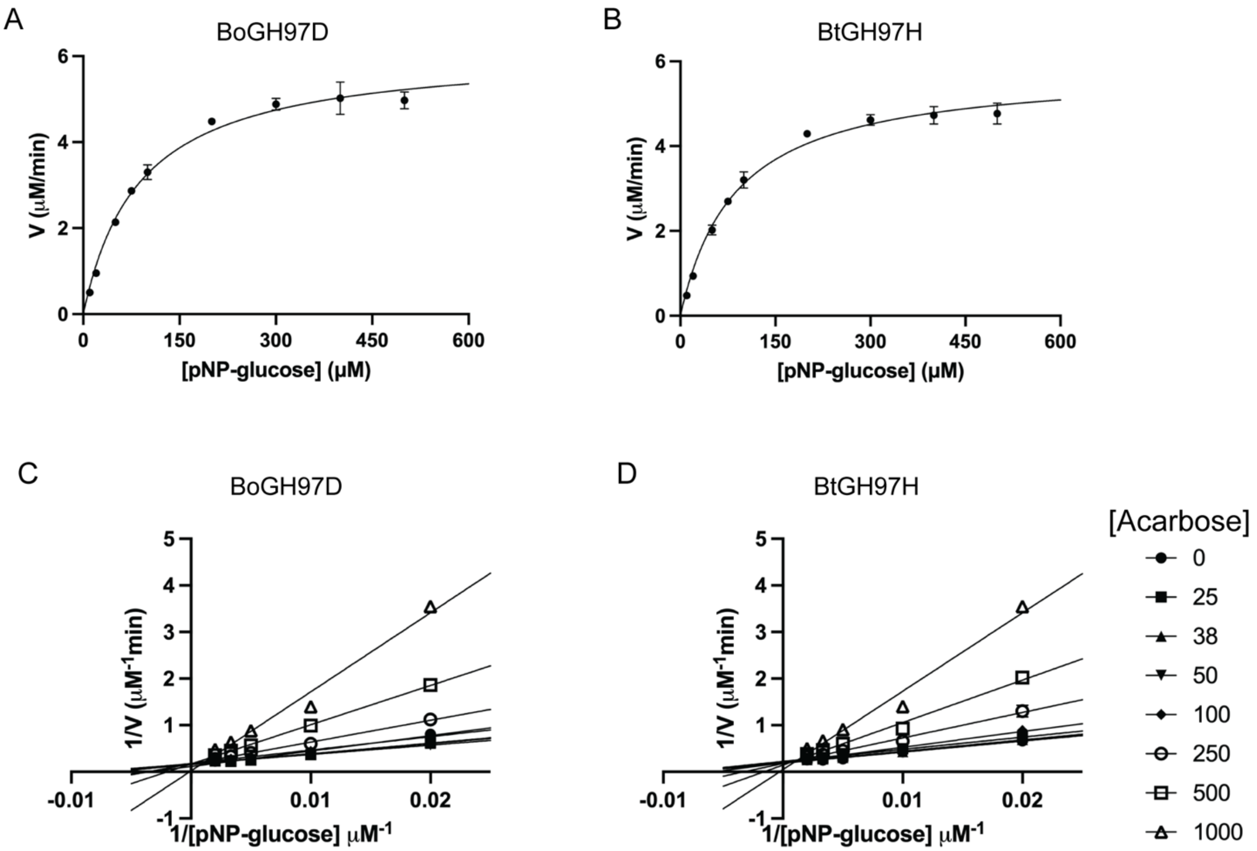


**Supplementary Figure 10. BoGH97D and BtGH97H are similarly inhibited by acarbose.** A,B) Enzymes were incubated for 10 min in various concentrations of pNP-glucose in duplicate. Initial rate as a function of substrate concentration is shown. C,D) Enzymes were pre-incubated for 10 min with acarbose before mixing with various pNP-glucose concentrations. Initial rates were recorded and used to make the double reciprocal Lineweaver-Burk plots shown. Michaelis-Menten and inhibition parameters can be found in Table 1. 2 nM of each enzyme was used for both experiments.

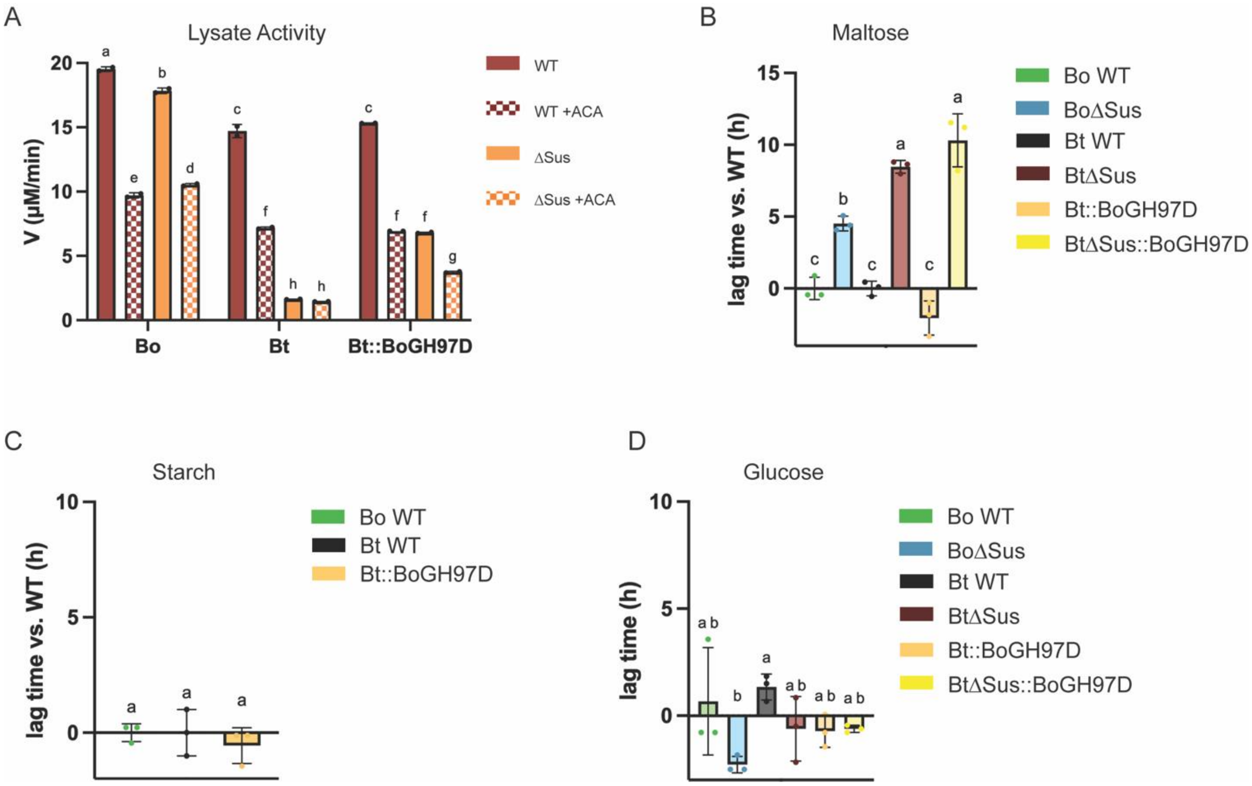


**Supplementary Figure 11. BoGH97D does not rescue the acarbose induced growth lag when expressed by Bt.** A) WT and ΔSus strains of Bo and Bt (with or without expressing BoGH97D from a constitutive promoter) were grown in minimal media (MM) + 5 mg/ml maltose to the same $OD_{600}$ and pelleted. Pellets were washed in PBS and cells were sonicated to release contents. Lysates were assayed in 1 mM pNP-Glc +/- 1 μM acarbose. B,C) Bacteria were pre-grown in MM with 5 mg/ml maltose and back diluted into MM + 2.5 mg/ml maltose or starch. Difference in time to $OD_{600}$ of 0.3 between the WT and mutant strains are graphed. D) Bacteria were pre-grown as in B/C and then grown in MM + 2.5 mg/ml glucose with or without 50 μM acarbose. Acarbose induced lag times are graphed. All statistical analyses were performed with a two-way ANOVA. Conditions with the same letter(s) were not significantly different from one another. A cutoff of $p \leq 0.05$ was used.

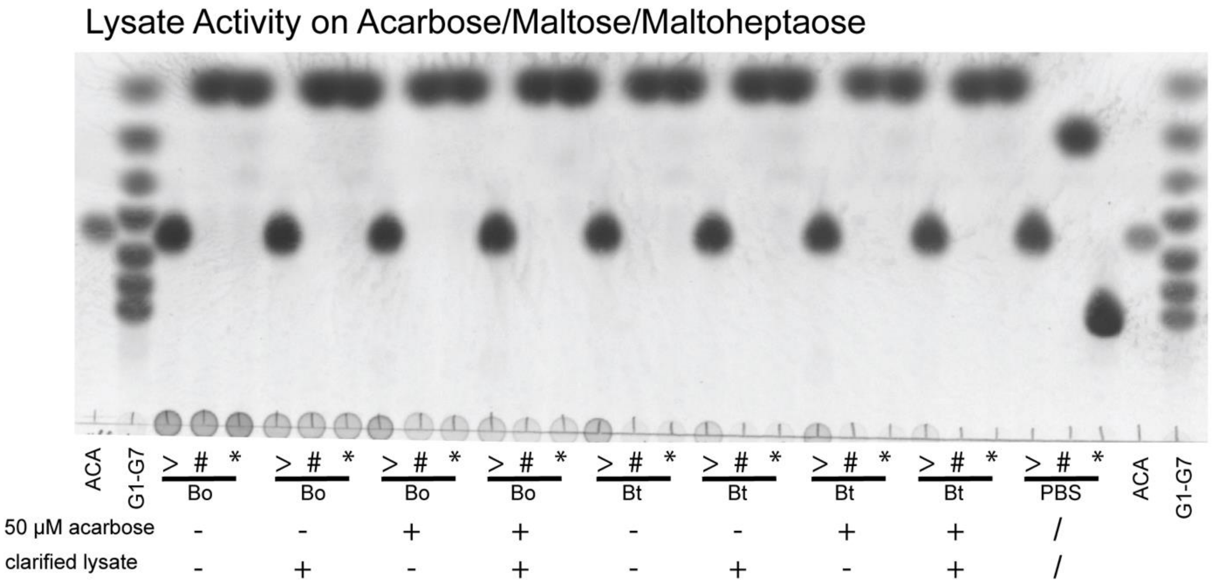


**Supplementary Figure 12.Lysates of Bo and Bt grown in maltose with acarbose do not break down acarbose.** Bo and Bt were grown in MM + 5 mg/ml maltose with or without 50 μM acarbose to an $OD_{600}$ of 0.7. Cells were pelleted and washed with PBS and sonicated to release cellular contents. Lysates and clarified lysates were incubated overnight at 37 °C with 5 mg/ml acarbose (>), maltose (#), or maltoheptaose(*). Clarified lysates were pelleted following cell sonication to remove insoluble components. 1 mg/ml acarbose and 1 mg/ml G1-G7 in PBS were run out on a thin layer chromatography (TLC) plate as reference points.
